# The SAGA acetyltransferase module is required for the maintenance of MAF and MYC oncogenic gene expression programs in multiple myeloma

**DOI:** 10.1101/2024.03.26.586811

**Authors:** Ying-Jiun C. Chen, Govinal Badiger Bhaskara, Yue Lu, Kevin Lin, Sharon Y. R. Dent

## Abstract

Despite recent advances in therapeutic treatments, multiple myeloma (MM) remains an incurable malignancy. Epigenetic factors contribute to the initiation, progression, relapse, and clonal heterogeneity in MM, but our knowledge on epigenetic mechanisms underlying MM development is far from complete. The SAGA complex serves as a coactivator in transcription and catalyzes acetylation and deubiquitylation. Analyses of datasets in the Cancer Dependency Map Project revealed many SAGA components are selective dependencies in MM. To define SAGA-specific functions, we focused on ADA2B, the only subunit in the lysine acetyltransferase (KAT) module that specifically functions in SAGA. Integration of RNA-seq, ATAC-seq, and CUT&RUN results identified pathways directly regulated by ADA2B include MTORC1 signaling, MYC, E2F, and MM-specific MAF oncogenic programs. We discovered that ADA2B is recruited to MAF and MYC gene targets, and that MAF shares a majority of its targets with MYC in MM cells. Furthermore, we found the SANT domain of ADA2B is required for interaction with both GCN5 and PCAF acetyltransferases, incorporation into SAGA, and ADA2B protein stability. Our findings uncover previously unknown SAGA KAT module-dependent mechanisms controlling MM cell growth, revealing a vulnerability that might be exploited for future development of MM therapy.

## Introduction

Multiple myeloma (MM) is a malignancy originating from antibody-producing plasma cells that reside within the bone marrow. Initiation and progression of this hematologic cancer are driven by genetic and epigenetic aberrations (Alzrigat et al. 2018b). Extensive genomic heterogeneity within individual tumors and across patients results in variations in response to treatment and overall survival rates. Primary genetic events that initiate MM formation include chromosomal hyperdiploidy and translocations that juxtapose *MMSET*, *CCND1*, *CCND3*, *MAFB* or *MAF* (also known as *c-MAF*) in proximity to the *IgH* enhancer (Bergsagel and Kuehl 2003). Secondary events that further promote MM progression include translocation of *MYC* (also known as *c-MYC*; t(8;14)), as well as gain– and loss-of-function mutations in oncogenes and tumor suppressor genes, respectively (Muylaert et al. 2022).

Translocation of *MAF* with *IgH* (t(14;16)) and other mechanisms give rise to MAF overexpression in approximately 50% of MM patients (Hurt et al. 2004; Zhan et al. 2006). MAF is a cell type-specific leucine zipper transcription factor (TF) of the AP-1 superfamily that regulates other oncogenes such as *CCND2* and *ITGB7* respectively encoding a cyclin involved in cell cycle and a transmembrane cell adhesion molecule. (Motohashi et al. 1997; Hurt et al. 2004; Alvarez-Benayas et al. 2021). Recently, MAF was defined as a pioneer TF that initiates the opening of closed chromatin for *de novo* transcriptional activation and myelomagenesis (Katsarou et al. 2023). In addition, MYC is a master oncogenic TF overexpressed in a majority of human cancers. MYC functionally collaborates with E2F1 TF to promote MM tumor growth through concerted regulation of the cell cycle and DNA replication (Zeller et al. 2006; Fulciniti et al. 2018).

Epigenetic dysregulation also contributes to oncogenesis, and the reversibility and plasticity of epigenetic modifications render them attractive targets for clinical therapy (Wu et al. 2020). For instance, global hypomethylation is associated with progression of the premalignant monoclonal gammopathy of undetermined significance (MGUS) phase to MM (Muylaert et al. 2022). DNA hypomethylating agents and inhibitors of histone deacetylases (HDACs) and histone modification readers have been approved for some hematologic malignancies, including a HDAC inhibitor (Panobinostat) and two acetylation reader inhibitors (JQ1, I-BET151) for the treatment of MM (Wu et al. 2020). However, MM remains largely incurable, with the occurrence of relapsed and refractory disease in many patients, underscoring the need for novel therapeutic approaches.

To date no lysine acetyltransferase (KAT) inhibitors have reached clinical trials for anti-cancer therapy. GCN5 (also known as KAT2A) was the first KAT to be linked to transcription (Brownell and Allis 1995; Brownell et al. 1996), and it has been shown to promote cancer growth and progression in MYC-induced B cell lymphoma, acute myeloid leukemia (AML), and solid cancers including hepatocellular carcinoma, colon, breast and non-small cell lung cancer (NSCLC) (Chen et al. 2013; Yin et al. 2015; Majaz et al. 2016; Tzelepis et al. 2016; Farria et al. 2019; Mustachio et al. 2019; Farria et al. 2020; Oh et al. 2020). GCN5 interacts with TFs such as MYC and E2F1 and serves as a coactivator of their gene targets through acetylation of histone H3 lysine 9 acetylation (H3K9ac) and non-histone proteins. Non-histone substrates of GCN5 include MYC and cell cycle regulators CDC6 and CDK9 (Patel et al. 2004; Downey 2021). GCN5 has a mammalian homologue termed PCAF (or KAT2B) and currently available inhibitors that target GCN5 also suppress PCAF activity. Nevertheless, the different phenotypes of their respective mutant mice indicate these homologues have both redundant and independent functions (Xu et al. 2000; Yamauchi et al. 2000), consistent with findings that these KATs have common as well as distinct substrates (Downey 2021).

In humans, GCN5 and PCAF are incorporated in a mutually exclusive manner in the KAT module of two different complexes, SAGA and ATAC, both of which act as coactivators in transcription (Grant et al. 1997; Suganuma et al. 2008; Wang et al. 2008; Chen and Dent 2021). Consequently, genetic or inhibitor studies on GCN5 functions cannot delineate whether the functions are executed through SAGA or ATAC complex. GCN5 and PCAF are required for H3K9ac in human cells, while ATAC houses a second KAT, ATAC2 (KAT14), that specifically targets histone H4 (Guelman et al. 2009). SAGA and ATAC have been shown to execute distinct biological functions in hematopoiesis and leukemia (Arede et al. 2022), therefore it is important to decipher their specific functions in different cancers for selective inhibition in therapy. The two KAT module are distinguished by the ADA2 component, with ADA2B specific to SAGA and ADA2A specific to ATAC (Kusch et al. 2003; Muratoglu et al. 2003). The ADA proteins are important cofactors for GCN5-mediated acetylation, as GCN5 alone displays very little activity on histone H3 and H4 peptides (Riss et al. 2015). Hence, paralogous ADA2B and ADA2A are ideal candidates to elucidate SAGA– or ATAC-specific KAT functions.

In this study, we demonstrate that in the context of MM, the SAGA complex is required for expression of MAF, MYC and E2F targets. We report for the first time a role for SAGA-specific ADA2B and KAT function in maintaining MM through sustaining both MYC and MAF protein levels and their downstream oncogenic gene expression programs. SAGA had not been previously associated with MAF in any context, and here we show that ADA2B also binds to MAF and MYC gene targets, further indicating ADA2B and SAGA serve as a coactivator for these TFs. We additionally demonstrate that the SANT domain of ADA2B is required for ADA2B protein stability and for interaction with GCN5 and PCAF, providing a vulnerability that might be exploited for development of future therapeutic interventions.

## Results

### SAGA-specific ADA2B is a dependency in multiple myeloma

To investigate SAGA-specific functions in cancer, we first centered on subunits that are exclusive to this complex, namely ADA2B, ADA1, SUPT3H, SUPT7L, SUPT20H, TAF5L, TAF6L, ATXN7, and USP22 (Figure 1A). Mining of the Cancer Dependency Map (DepMap) datasets (Pacini et al. 2021) uncovered a dependence of lymphoid and myeloid cells on *ADA2B* (Figure 1B), consistent with previous studies by our group and others reporting GCN5 is required for lymphoma and AML cell growth (Tzelepis et al. 2016; Farria et al. 2019; Farria et al. 2020). Moreover, we found that *ADA2B* is highly expressed in plasma cell-derived MM cells and is exceptionally crucial for their growth and survival as compared to other blood cancers and solid tumor lineages (Figure 1B).

**Figure 1.**
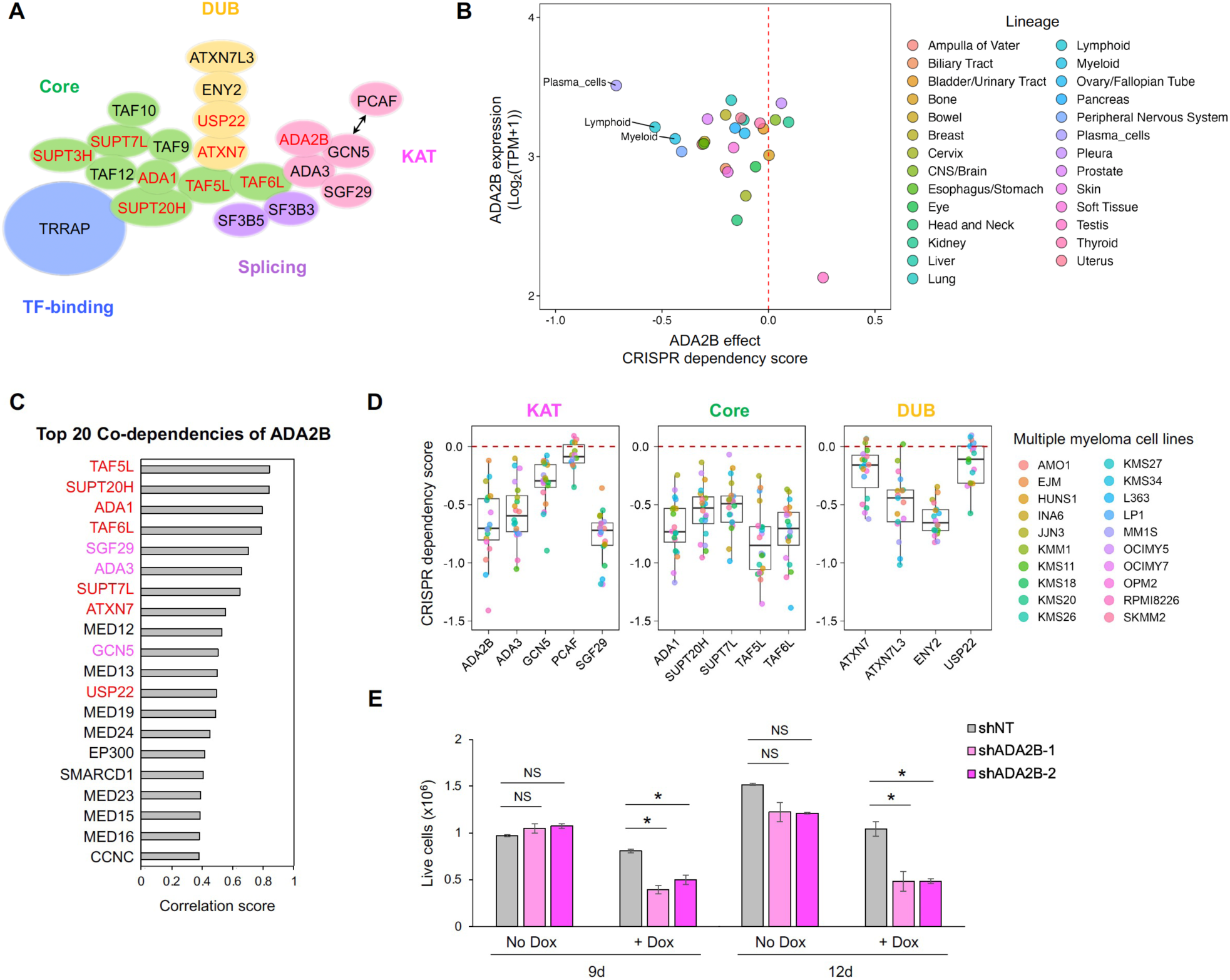
SAGA-specific ADA2B is a dependency in MM. **A)** Components of the SAGA complex. Lysine acetyltransferase (KAT), deubiquitinase (DUB), splicing, core, and transcription factor (TF)-binding modules are shown in pink, yellow, purple, green, and blue, respectively. Components specific to SAGA are indicated in red. Relative positions of components are illustrated based on the previously reported cryogenic-electron microscopy structure of human SAGA (Herbst et al. 2021). **B)** The mean values of dependency scores and expression of ADA2B across cancer cell lines derived from various lineages found in CRISPR DepMap and Sanger (Score) databases (DepMap 23Q2+ Score, Chronos). **C)** Top twenty co-dependencies of ADA2B across the same cancer cell lines as in panel B. Components specific to SAGA complex are indicated in red. SAGA KAT module components are indicated in pink. **D)** Dependency scores of SAGA components in KAT, core, and DUB modules for multiple myeloma cell lines. **E)** Cell viability assay of Dox-inducible shRNA lines. A one-way Analysis of Variance (ANOVA) was performed to analyze the differences among group means, followed by the Tukey HSD post hoc test to determine whether the mean difference between specific pairs of group are statistically significant. * p < 0.05, not significant (NS) p > 0.05.

Further supporting a role for the SAGA complex in cancer cell growth, the top ten co-dependencies (genes with similar dependency scores and thus likely confer similar functions) of *ADA2B* in the DepMap datasets included nine genes encoding other SAGA subunits. Six of these encoded components function exclusively in the SAGA complex and three function in the KAT module, consistent with the known structure and function of the SAGA complex (Figures 1C and 1D). The positive correlation of the cancer dependency scores of *ADA2B* and *TAF5L*, in contrast to the negative correlation of the dependency scores of *ADA2B* and ATAC-specific *ADA2A*, further indicate that ADA2B serves as a proxy for the SAGA complex (Figure S1B). Interestingly, seven genes encoding components of the Mediator complex also scored in the top twenty ADA2B co-dependencies, suggesting a cooperation between Mediator and SAGA in cancer cells (Figure 1C).

To further determine which SAGA functions are most critical for survival of MM cells, we compared dependency scores of SAGA subunit genes from each module. MM cell lines are highly dependent on genes from the KAT, core, TF-binding and splicing modules, as indicated by their low dependency scores (Figures 1D and S1C). However of all the SAGA-specific components, only *ADA2B* and core subunit genes that are among the top *ADA2B* co-dependencies (*ADA1*, *SUPT7L*, *SUPT20H*, *TAF5L*, *TAF6L*) are genetic vulnerabilities in MM cells (Figures 1A, 1C and 1D). In contrast, SAGA-specific and deubiquitinase (DUB) module genes *ATXN7* and *USP22* are non-essential. Paralogous *GCN5* and *PCAF* are individually not essential for MM cell survival, likely due to redundant functions of these KATs. The strong positive correlation of *GCN5* and *PCAF* dependency scores in MM cell lines further supports these homologues execute similar functions in this cell type (Figure S1B).

To complement above findings in cultured cells, we also compared expression of SAGA subunit genes in MM patients to their expression in individuals with pre-malignant MGUS, asymptomatic smoldering myeloma, or healthy plasma cells using publicly available microarray datasets (Hanamura et al. 2006; Zhan et al. 2007). Nearly half of SAGA subunit genes, including *GCN5*, are overexpressed in a vast majority of MM patients. The remainder SAGA subunit genes, including *ADA2B*, are overexpressed in approximately 50% of MM cases (Figure S1A).

As ADA2B is the only SAGA-specific MM dependency functioning in the therapeutically more tractable enzymatic modules, we focused on ADA2B for further studies. To validate the essentiality of ADA2B to MM viability, we generated doxycycline (Dox)-inducible knockdown (KD) of ADA2B in the MM.1S cell line. Transcriptome profiling by others indicates MM.1S is a top-ranked cell line highly representative of MM patient tumors (Sarin et al. 2020). Consistent with data from DepMap, we observed reduced cell viability upon Dox-induced ADA2B depletion (Figure 1E). This phenotype was also reproduced in our constitutive CRISPR-mediated knockout (KO) lines of ADA2B (Figure S1D). Reduced levels of ADA2B protein were confirmed at 3 days and 9 days of Dox treatment by immunoblot, with a greater reduction observed upon longer treatment time (Figure 2A). As expected, since ADA2B is a required component for SAGA KAT activity, we observed decreased global H3K9ac levels upon ADA2B depletion. Surprisingly, both GCN5 and PCAF levels were also reduced upon ADA2B loss, with the impact on GCN5 levels occurring earlier than PCAF.

**Figure 2.**
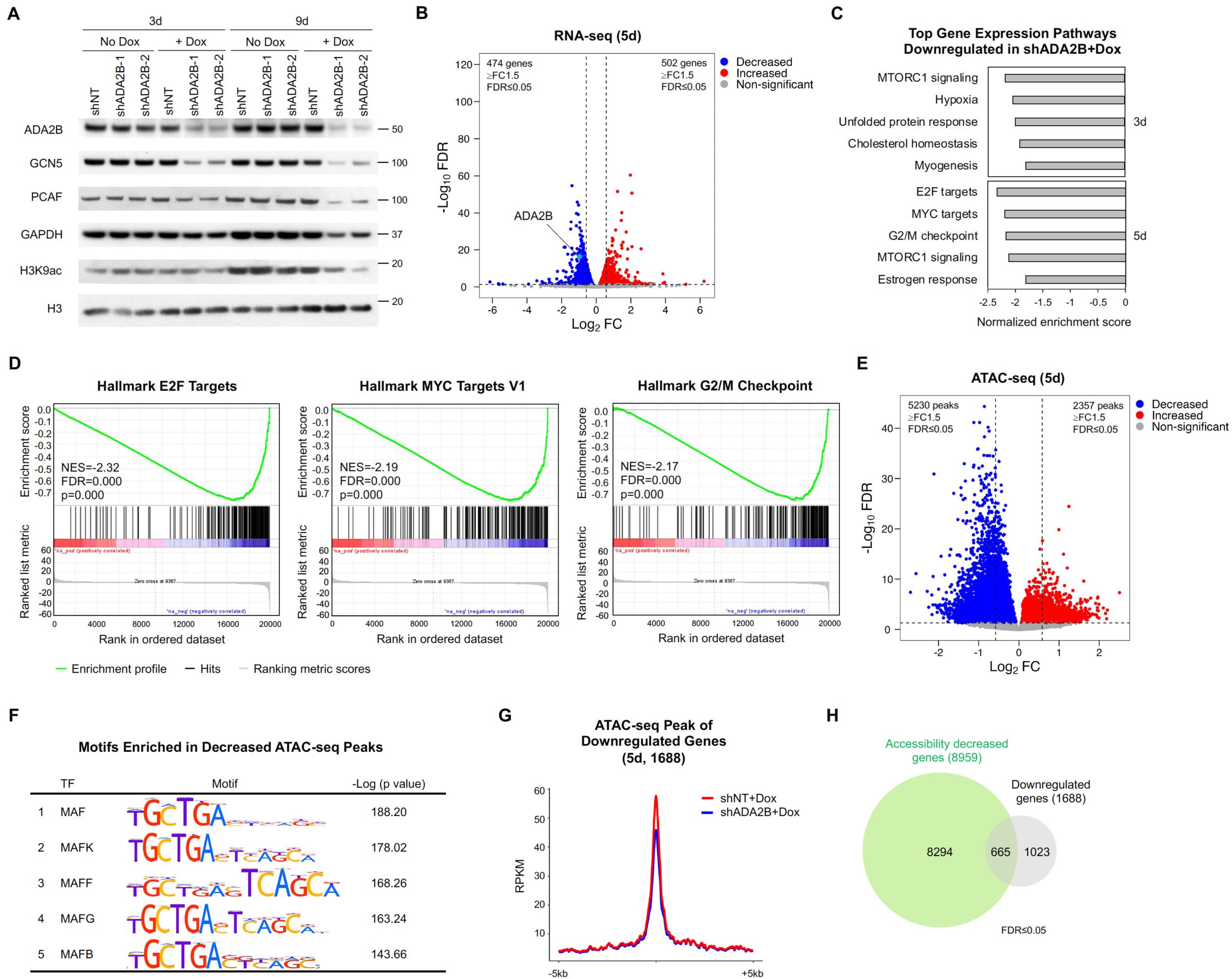
ADA2B is required for maintenance of oncogenic gene expression programs. **A)** Immunoblots showing levels of ADA2B, GCN5, PCAF, and H3K9ac in shNT and shADA2B cells with and without Dox treatment. Histone H3 is included as a loading control. **B)** Volcano plot of gene expression changes detected by RNA-seq in shADA2B cells relative to shNT cells after 5 days of Dox treatment. Genes with statistically significant differential expression with fold change (FC) ≥ 1.5 include 474 downregulated genes and 502 upregulated genes. **C)** Top five gene expression pathways identified by GSEA that were enriched in downregulated genes in shADA2B cells relative to shNT cells after 3 and 5 days of Dox treatment. **D)** Enrichment plots for E2F targets, MYC targets v1, and G2/M checkpoint hallmarks. **E)** Volcano plot of chromatin accessibility changes detected by ATAC-seq in shADA2B cells relative to shNT cells after 5 days of Dox treatment. Statistically significant differential peaks with FC ≥ 1.5 include 5230 decreased peaks and 2357 increased peaks. **F)** Top five TF motifs enriched in decreased ATAC-seq peaks in shADA2B cells relative to shNT cells after 5 days of Dox treatment. **G)** Average RPKM (reads per kilobase per million) normalized signal of ATAC-seq peaks associated to all (1688) downregulated genes in shADA2B cells relative to shNT cells after 5 days of Dox treatment. **H)** Venn diagram depicting the overlap between downregulated genes (1688 genes) and genes with decreased accessibility (8959 genes) in shADA2B cells relative to shNT cells after 5 days of Dox treatment.

### ADA2B is required for maintenance of oncogenic gene expression programs

To identify the gene expression pathways regulated by ADA2B in MM, we performed total RNA sequencing (RNA-seq) on the Dox-inducible KD MM.1S lines treated for 3 and 5 days in parallel with a Non-targeting (NT) shRNA line and untreated cells as controls (Figure S2A). To eliminate non-specific effects from Dox treatment, we compared the Dox-induced KD line to Dox-treated NT cells. At 3 days, 3-fold more genes were markedly downregulated (533 genes FC ≥ 1.5, FDR ≤ 0.05) than upregulated (177 genes FC ≥ 1.5, FDR ≤ 0.05) in the ADA2B-depleted cells relative to treated NT line (Figures S2A and S2B), consistent with the SAGA functions as a transcriptional coactivator. Downregulation persisted to the later time point for 50% of all genes with decreased expression (Figure S3A, Table S1). At 5 days, markedly upregulated genes (502 genes FC ≥ 1.5, FDR ≤ 0.05) increased to a similar quantity as downregulated genes (474 genes FC ≥ 1.5, FDR ≤ 0.05), which is likely due to secondary effects (Figure 2B).

Gene Set Enrichment Analysis (GSEA) identified the top enriched pathways in downregulated genes as MTORC1 signaling at 3 days, and E2F, MYC gene targets, and G2/M checkpoint genes at 5 days (Figures 2C, 2D and S2C). These findings are consistent with previous studies establishing GCN5 as a coactivator for E2F and MYC gene targets in other settings, including embryonic stem cells (ESCs) and in cancers such as Burkitt’s lymphoma and NSCLC (Hirsch et al. 2015; Farria et al. 2019; Mustachio et al. 2019; Farria et al. 2020). Consistent with our cell viability phenotype, the downregulated MTORC1 signaling, E2F and MYC pathways are well-known for promoting MM development (Chesi et al. 2008; Li et al. 2014; Fulciniti et al. 2018). Also in accordance with the cellular phenotype, p53 tumor suppressor pathway genes, which play a major role in the inhibition of cancer cell growth (Hernández Borrero and El-Deiry 2021), were enriched in the upregulated genes at 5 days of treatment (Figure S2D).

The global decrease in H3K9ac observed upon ADA2B depletion suggests that chromatin accessibility might be affected. To address this question, we performed Assay for Transposase-Accessible Chromatin with sequencing (ATAC-seq) using the same set of samples as for RNA-seq. At both 3– and 5-day time points, we observed a much larger number of decreased ATAC-seq peaks than increased peaks (Figures 2E and S2E), indicating that ADA2B depletion is associated mostly with decreased chromatin accessibility, in accordance with the long-standing link between histone acetylation and open chromatin (Chen et al. 2022). We also observed a major overlap of genes with decreased accessibility at the two time points (Figure S3B, Table S1).

Motif analysis of decreased ATAC-seq peaks identified the top five enriched binding sites as motifs of the MAF family TFs (Figure 2F). Accessibility of downregulated genes upon ADA2B depletion was decreased on average (Figures 2G and S2F), and approximately one-third of genes with lower expression also had decreased accessibility (Figures 2H and S2G), including oncogenes *SH3RF2*, *PCAT1* and *MITF* that are ectopically expressed in human cancers (Figures S3C).

### Cell cycle related genes are ADA2B core targets

We next determined genomic sites bound by ADA2B in order to define its direct targets in MM. Since ChIP-grade ADA2B antibodies are not readily available, we created MM.1S cell lines expressing epitope-tagged ADA2B (FLAG-3xHA-ADA2B) or the tags alone (FLAG-3XHA) as a control for off-target antibody binding. We successfully performed Cleavage Under Targets and Release Using Nuclease assay (CUT&RUN) for ADA2B binding in parallel with H3K9ac profiling (Figure 3A). Our results identified 3,344 ADA2B binding sites that were statistically significant in at least two replicates (Figure 3B). Consistent with previously described SAGA binding sites in other contexts (Arede et al. 2022), ADA2B and H3K9ac predominantly locate to accessible chromatin regions determined by ATAC-seq and promoter regions in the genome (Figures 3A and 3B). To analyze the functional enrichments of the genes that are bound by ADA2B at their putative promoters, we performed Ingenuity Pathway Analysis (IPA). The top enriched pathways include cell cycle-related mitotic prophase and sirtuin signaling pathways (Figure S4A). The top upstream regulators identified include HNF4A, MYC, and E2F family members, E2F4 and E2F1. These findings correspond to the enrichment of MYC and E2F targets in genes downregulated upon ADA2B depletion identified by RNA-seq (Figure S4B).

**Figure 3.**
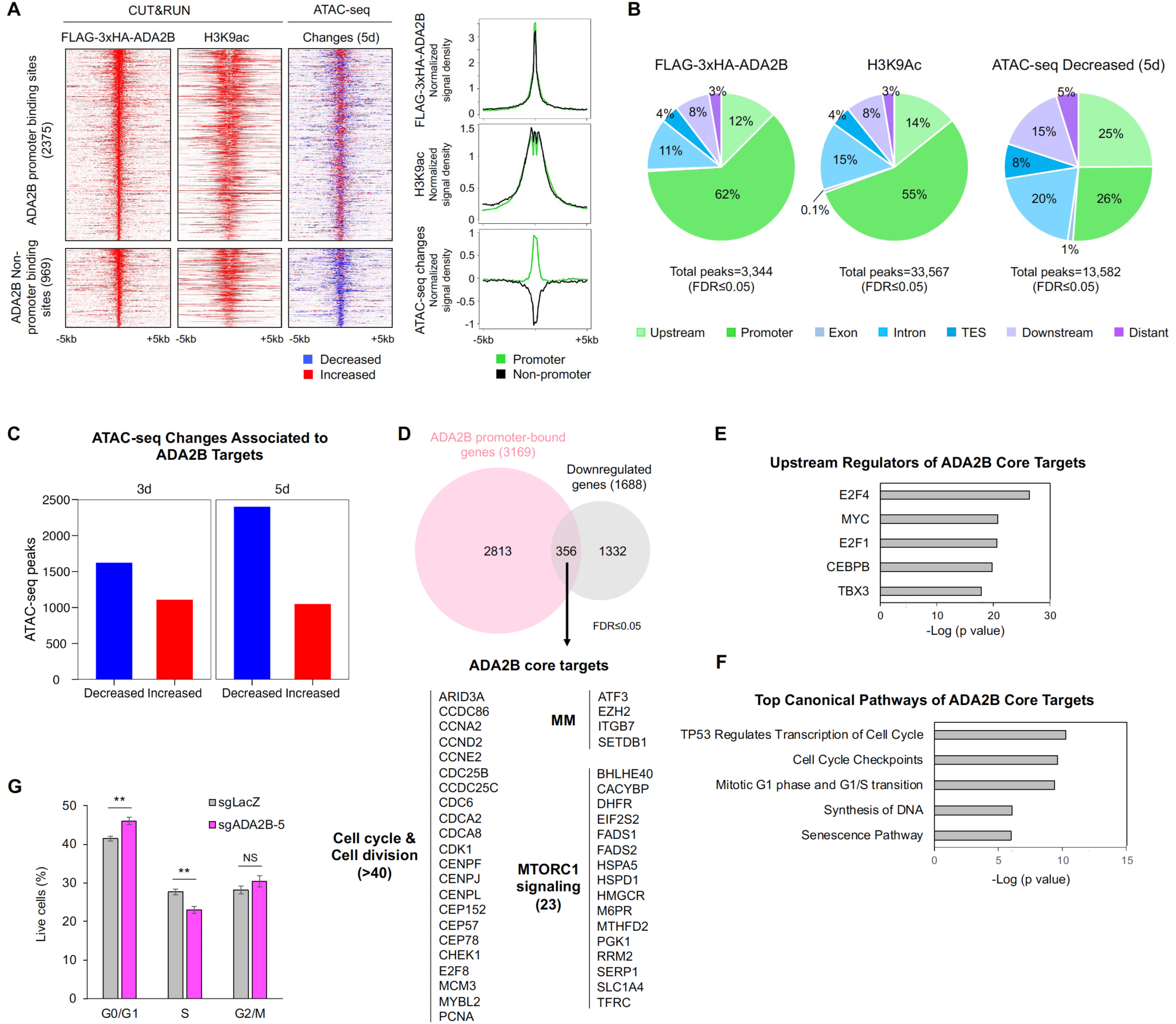
ADA2B is recruited to promoters of genes involved in tumorigenesis. **A)** Heatmaps of FLAG-3xHA-ADA2B peaks, H3K9ac peaks in cells expressing FLAG-3xHA, ATAC-seq peaks in Dox-treated shNT and shADA2B cells, and changes of ATAC-seq peak signals in shADA2B cells relative to shNT cells after 5 days of Dox treatment. Decreased peaks are shown in blue; increased peaks are shown in red. Regions are ranked based on ADA2B signal. FLAG-3xHA-ADA2B signals are normalized to IgG signals. ChIP-seq tracks of MAF, MYC, and Pol II represent FC over input. Metaplots show average normalized signal density for FLAG-3xHA-ADA2B, H3K9ac, and changes in ATAC-seq peaks. **B)** Pie chart depicting genomic distribution of FLAG-3xHA-ADA2B, H3K9ac in cells expressing FLAG-3xHA, and decreased ATAC-seq peaks in shADA2B cells relative to shNT cells after 5 days of Dox treatment. **C)** Bar chart showing quantity of decreased and increased ATAC-seq peaks associated to ADA2B-bound genes in shADA2B cells relative to shNT cells after 3 and 5 days of Dox treatment. **D)** Venn diagram depicting the overlap between ADA2B promoter-bound genes (3169 genes) and downregulated genes (1688 genes) in shADA2B cells relative to shNT cells after 5 days of Dox treatment, identifying 356 ADA2B core targets. ADA2B core targets include more than 40 genes involved in cell cycle and cell division, 23 genes in MTORC1 signaling, and known regulators of MM biology. **E)** Top five upstream regulators of ADA2B core targets identified by IPA. **F)** Top five canonical pathways of ADA2B core targets identified by IPA. **G)** Cell cycle analysis of sgADA2B and control sgLacZ cells. The percentage of events in each cell cycle stage for three biological replicates is graphed. A one-way Analysis of Variance (ANOVA) was performed to analyze the differences among group means, followed by the Tukey HSD post hoc test to determine whether the mean difference between specific pairs of group are statistically significant. ** p < 0.01, * p < 0.05, not significant (NS) p > 0.05.

Genes bound by ADA2B were often associated with decreased accessibility in ADA2B KD cells, especially at the later time point (Figure 3C). Interestingly, we observed a pattern of lower accessibility upon ADA2B depletion at non-promoter binding sites compared to promoter binding sites of ADA2B (Figure 3A). Indeed, only a quarter (26%) of decreased ATAC-seq peaks was located at promoter regions (Figure 3B).

To identify ADA2B core targets, we integrated genes bound by ADA2B at their putative promoters with genes downregulated upon ADA2B depletion. We identified 214 and 356 core targets of ADA2B in 3-day and 5-day ADA2B KD cells, respectively (Figures 3D and S4C, Table S2). Over 20% of downregulated genes in 5-day ADA2B KD cells were directly targeted by ADA2B. IPA analysis revealed ADA2B core targets are again enriched with genes involved in cell cycle, cell division, and MTORC1 signaling, with top upstream regulators identified as MYC, E2F4 and E2F1 (Figures 3E and 3F). Cell cycle analysis of ADA2B KO cells by flow cytometry showed increased accumulation at the G0/G1 stage (Figure 3G), corresponding to the enrichment of G1/S transition related genes among ADA2B core targets. Additionally, many MTORC1 signaling genes directly targeted by ADA2B facilitate cancer growth and progression (Figure 3D) (He et al. 2019b; Kiss et al. 2020; Mo et al. 2020; Zhao et al. 2020; Shan et al. 2022; Feng et al. 2023). Known regulators of MM biology, such as *EZH2*, *ITGB7*, and *SETDB1*, were also among the ADA2B core targets (Neri et al. 2011; Alzrigat et al. 2018a; Qian et al. 2023).

### ADA2B loss impacts MAF protein levels and MAF-driven oncogenic gene expression programs

Given the enrichment of MYC and E2F1 gene targets among the ADA2B core targets, we asked if MYC and E2F1 expression are altered by ADA2B depletion. MYC is a known substrate of both GCN5 and PCAF, and acetylation stabilizes MYC protein levels (Patel et al. 2004). E2F1 is also targeted by several KATs including PCAF (Ianari et al. 2004). Indeed in ADA2B KD cells, we observed decreased levels of MYC and E2F1 proteins (Figure 4A). While some decrease in E2F1 RNA levels was observed, MYC RNA levels were unaffected or slightly increased at the later time point, indicating the reduced MYC protein expression is attributed to post-translational regulation, whereas reduced E2F1 expression may be the result of both transcriptional and post-translational regulation (Figure S5A).

**Figure 4.**
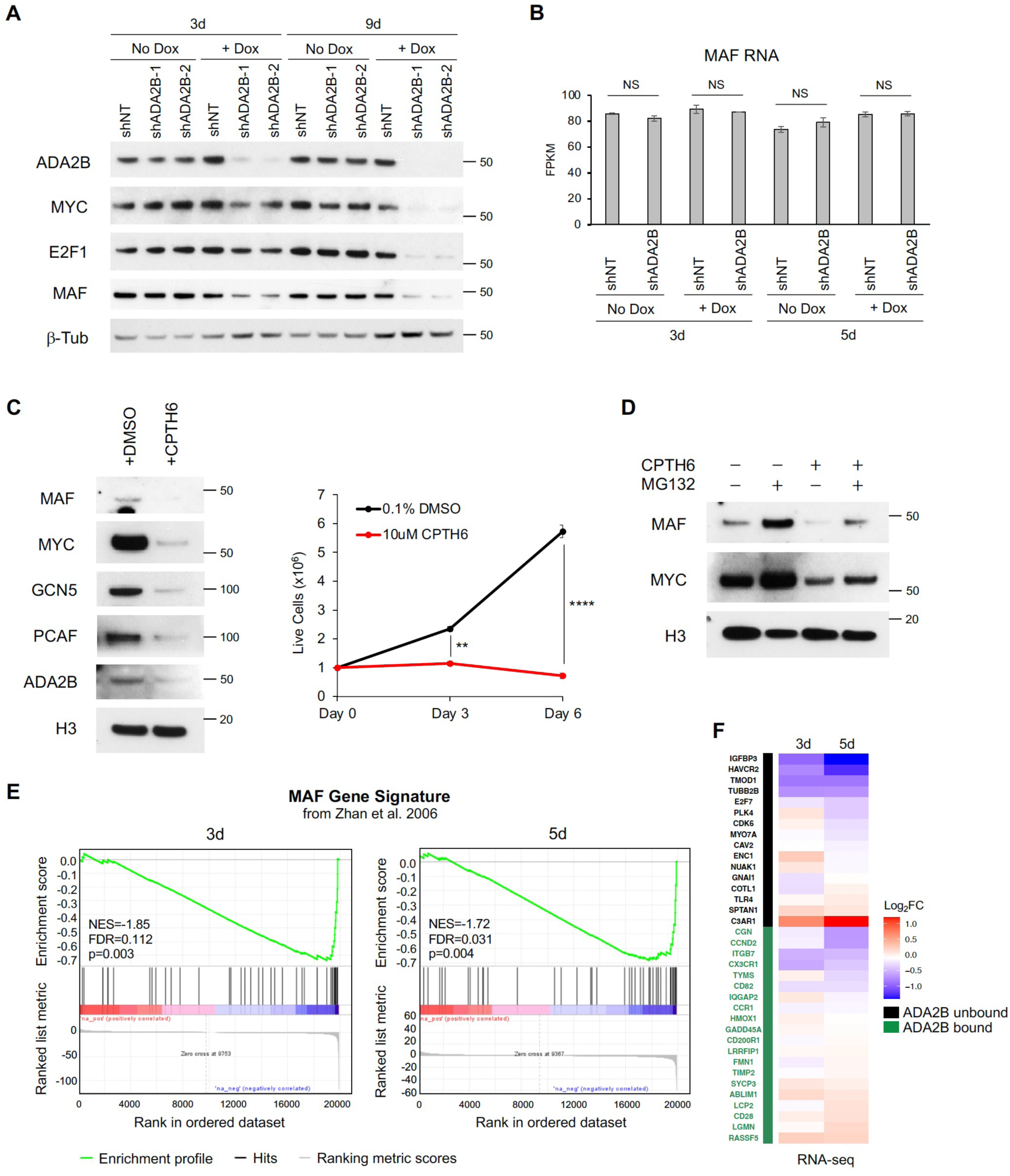
ADA2B promotes MAF protein levels and MAF expression program. **A)** Immunoblots showing levels of MYC, E2F1, and MAF in shNT and shADA2B cells with and without Dox treatment. b-Tub is included as a loading control. **B)** Transcript levels in FPKM (fragments per kilobase per million) of MAF in shNT and shADA2B cells with and without Dox treatment as determined by RNA-seq. **C)** Effects of CPTH6 treatment on MM.1S cells. Left: immunoblots showing MAF, MYC, GCN5, PCAF, ADA2B levels in MM.1S cells treated with 10 µM CPTH6 for 3 days in comparison to DMSO vehicle control. Histone H3 is included as a loading control. Right: Cell viability assay of MM.1S cells treated with 10 µM CPTH6. A one-way Analysis of Variance (ANOVA) was performed to analyze the differences among group means, followed by the Tukey HSD post hoc test to determine whether the mean difference between specific pairs of group are statistically significant. **** p < 0.0001, *** p < 0.001, ** p < 0.01, * p < 0.05, not significant (NS) p > 0.05. **D)** Immunoblots of MM.1S cells with or without 10 µM CPTH6 treatment for 3 days and subsequent treatment of DMSO or 10 µM MG132 for 5 hours. Histone H3 is included as a loading control. **E)** GSEA enrichment plots demonstrating enrichment of MAF gene signature from Zhan et al. 2006 (Zhan multiple myeloma MF up) in the genome-wide expression changes induced by Dox in shADA2B cells compared to shNT cells for 3 and 5 days. **F)** Heatmap showing expression of selected MAF core targets in shADA2B cells compared to shNT cells after 3 and 5 days of Dox treatment. Genes directly bound by ADA2B as determined by our CUT&RUN data are highlighted in green.

We next questioned whether ADA2B might regulate other oncogenic TFs that are specifically expressed in MM, such as IRF4 and MAF. We found that MAF protein levels are strongly decreased in ADA2B-depleted MM cells (Figure 4A). IRF4 levels were also noticeably reduced (Figure S5B). Protein abundance of DP1, partner of E2F1 required for MM cell proliferation (Fulciniti et al. 2018), was not consistently affected (Figure S5B). Transcript levels of MAF were unchanged (Figure 4B), suggesting that ADA2B and SAGA may regulate MAF levels through post-translational acetylation. To test this hypothesis, we treated MM.1S cells with CPTH6, an inhibitor that targets the KAT activity of GCN5 and PCAF (Trisciuoglio et al. 2012). CPTH6-treated MM.1S cells displayed severe growth arrest (Figure 4C), indicating MM cells are highly dependent on the KAT activity of GCN5/PCAF for growth. Immunoblot analysis again revealed a clear reduction of MAF as well as MYC and global H3K9ac levels (Figures 4C and S5C). CPTH6 treatment also resulted in reduction of both GCN5 and PCAF. These changes are consistent with previous findings demonstrating that PCAF auto-acetylates and that *GCN5* gene expression is directly promoted by MYC (Knoepfler et al. 2006; Yin et al. 2015; Downey 2021). Interestingly, ADA2B levels also decreased with CPTH6 treatment, suggesting the inhibitor impacts the stability of the SAGA KAT module.

To determine if MAF protein levels are regulated by proteasomal degradation upon inhibition of GCN5/PCAF activity, we subjected MM.1S cells to combined treatment of CPTH6 with the proteasome inhibitor MG132. Addition of MG132 increased MAF and MYC abundance in both CPTH6 treated and untreated cells (Figure 4D). Altogether, these results suggest that acetylation by the KAT module of the SAGA complex is directly or indirectly required for maintenance of MAF protein levels in MM cells.

An important role of ADA2B and SAGA in maintaining MAF functions is further demonstrated by downregulation upon ADA2B depletion of the MAF gene signature previously defined in MM patients (Zhan et al. 2006) (Figure 4E). Moreover, 45% of MAF core gene targets involved in cell cycle, cell migration and adhesion, p53 signaling, actin binding, cytokine and chemokine pathways (pathways that involved 5 or more MAF core targets; (Katsarou et al. 2023)) were decreased in expression upon ADA2B depletion (Figure 4F). Taken together, our results indicate ADA2B loss negatively impacts MAF protein abundance and MAF-driven oncogenic expression programs.

### ADA2B is recruited to MAF and MYC gene targets

Our CUT&RUN results showed that ADA2B directly binds to greater than 50% of the examined MAF core gene targets (Figure 4F). We further expanded our analysis to all MAF targets, utilizing published datasets of chromatin immunoprecipitation with sequencing (ChIP-seq) that were performed in the MM.1S cell line (Katsarou et al. 2023). MYC genomic occupancy in MM.1S was additionally included in our analyses to define co-targets of ADA2B and MYC in MM (Lin et al. 2012). Most ADA2B-bound genomic sites (2798 of 2985) were shared with MYC, and over 50% (1492) were additionally shared with MAF (Figures 5A and 5B). Surprisingly, our results revealed a large overlap between MYC and MAF targets (Figures 5B and S6A). Of the annotated 2317 shared target genes, 1687 genes had colocalized binding sites of all 3 regulators, suggesting that they may work in concert (Figure S6A). IPA analysis showed co-bound sites were located at genes enriched with functions in the deubiquitylation pathway and mitotic prophase (Figure S6B).

**Figure 5.**
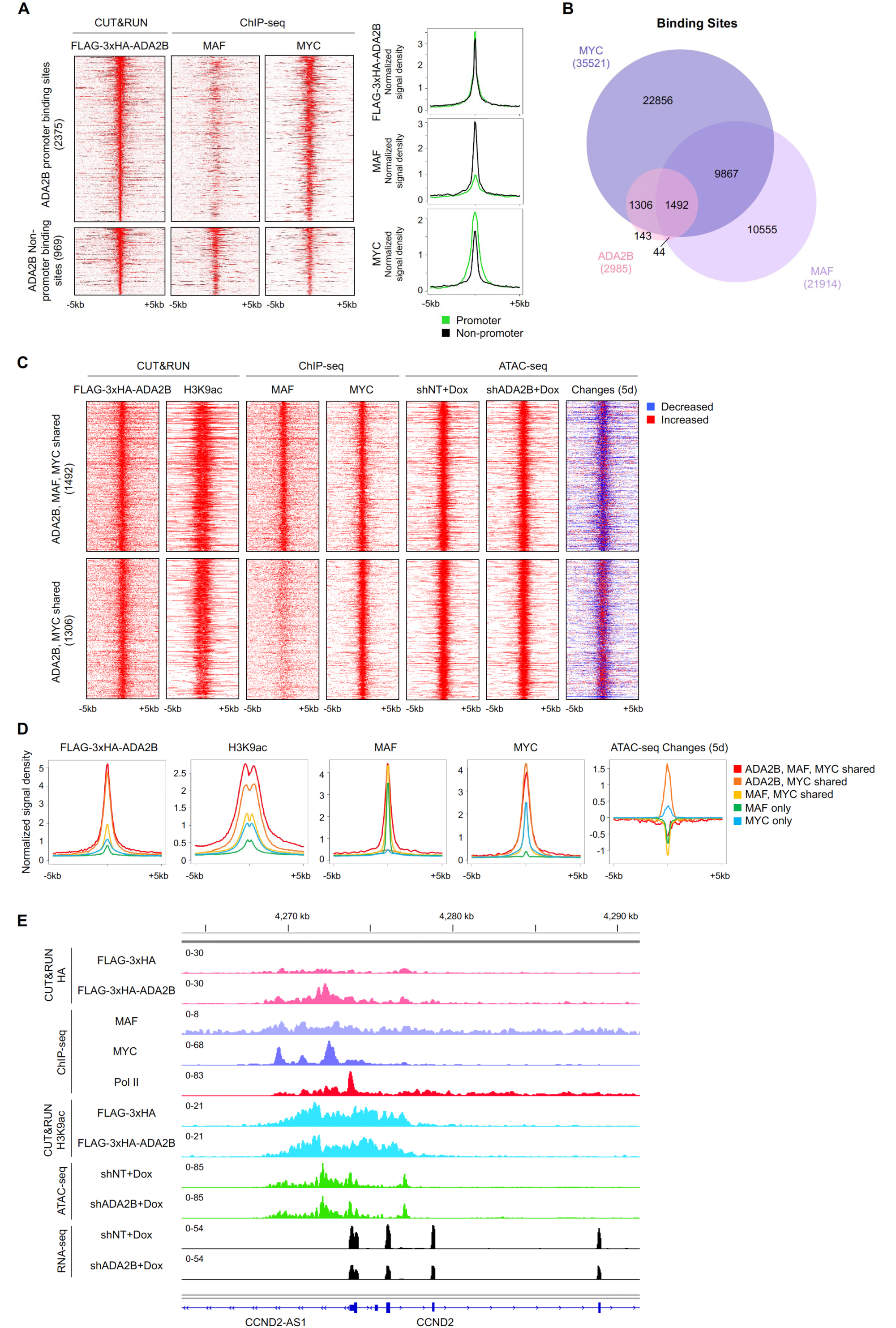
ADA2B directly binds to MAF and MYC targets. **A)** Heatmaps of FLAG-3xHA-ADA2B, MAF, and MYC binding peaks from our CUT&RUN analysis for ADA2B and from published ChIP-seq datasets for MAF and MYC (Lin et al. 2012; Katsarou et al. 2023). Regions are ranked based on ADA2B signal. FLAG-3xHA-ADA2B signals are normalized to IgG signals. ChIP-seq tracks of MAF, MYC, and Pol II represent FC over input. Metaplots show average normalized signal density for FLAG-3xHA-ADA2B, MAF, and MYC peaks. **B)** Venn diagram depicting the overlaps between ADA2B, MAF, and MYC binding sites (peaks). **C)** Heatmaps of FLAG-3xHA-ADA2B, MAF, MYC, and ATAC-seq peaks at ADA2B, MAF, and MYC shared binding sites and ADA2B and MYC shared binding sites. FLAG-3xHA-ADA2B signals are normalized to IgG signals. ChIP-seq tracks of MAF, MYC, and Pol II represent FC over input. Changes of ATAC-seq peak signals in shADA2B cells relative to shNT cells after 5 days of Dox treatment are also shown, with decreased peaks are in blue and increased peaks in red. **D)** Metaplots of FLAG-3xHA-ADA2B, MAF, MYC, and changes of ATAC-seq peaks shown in panel C and Supplemental Figure S6D. **E)** Binding patterns of FLAG-3xHA-ADA2B, MAF, MYC, Pol II, and H3K9ac at CCND2 locus. Peak signals from ATAC-seq and RNA-seq in shADA2B and shNT cells after 5 days of Dox treatment are also shown. ChIP-seq tracks of MAF, MYC, and Pol II represent FC over input.

MAF binding at ADA2B-bound sites is more robust at non-promoter regions compared to promoters (Figure 5A), consistent with previous work reporting MAF binding sites are mainly located at intergenic and intronic regions in the genome (Katsarou et al. 2023), whereas a reversed pattern is observed for MYC. Heatmap analysis of occupancy at shared and non-shared target regions revealed ADA2B-bound sites are enriched with H3K9ac and are located at more accessible chromatin (Figures 5C, 5D and S6D). Binding of MYC is more robust at ADA2B-bound sites than at MAF and/or MYC targets not bound by ADA2B, consistent with the higher accessibility. We also observed the pattern of lower accessibility upon ADA2B depletion at MAF-bound sites compared to MAF-unbound sites, which corresponds to the enrichment of MAF motif in decreased ATAC-seq peaks (Figures 2F and 5D).

The landscape of bound factors at *CCND2*, an oncogene important for cell cycle regulation, a well-known target of MAF, and an ADA2B core target, illustrates ADA2B, MAF, and MYC bind to the promoter region nearby the TSS and sites of RNA polymerase II (Pol II) occupancy (Figures 3D and 5E). The binding sites of these three regulators also colocalize with H3K9ac marks and the open chromatin region. Another core gene target of both ADA2B and MAF, *ITGB7*, which regulates cell adhesion, migration, and invasion in MM, is also bound by MYC (Figure S6C) (Neri et al. 2011). ADA2B, MAF, and MYC strongly colocalize with each other and with Pol II at *ITGB7*. Taken together, our data demonstrate that ADA2B is recruited to MAF and MYC gene targets, and that MAF shares a majority of its targets with MYC in MM cells.

### The SANT domain of ADA2B is required for GCN5/PCAF interaction, SAGA incorporation, and ADA2B protein stability

No small molecule inhibitors to directly suppress ADA2B function are currently available. Therefore we next sought to define which region of ADA2B is most critical for its functions in MM cell survival. ADA2B contains three annotated domains, a N-terminal ZZ-type zinc finger, a SANT domain, and a C-terminal SWIRM domain (Figures 6A and S7A) (Gamper et al. 2009). We utilized the ProTiler software to fine-map regions sensitive to CRISPR-mediated KO (He et al. 2019a). The predicted essential region matched precisely to the SANT domain (Figure 6A). The SANT domain is required for interaction with GCN5 and prevents the dissociation of acetyl-coenzyme A from the GCN5 KAT domain (Gamper et al. 2009; Sun et al. 2018). To further investigate the function of the SANT domain in MM, we generated constructs of ADA2B bearing point mutations of three conserved amino acid residues that are predicted to form the hydrophobic core of the domain (Boyer et al. 2004; Jumper et al. 2021; Varadi et al. 2021), either mutating the residues to alanine or to another bulky residue, phenylalanine (W70A, W90A, Y110A and W70F, W90F, Y110F; Figures 6B, 6C, S7A and S7B). The transcript levels of exogenous, mutated ADA2B (FLAG-3xHA-ADA2B^W70A W90A Y110A^ and FLAG-3xHA-ADA2B^W70F^ ^W90F^ ^Y110F^) were similar to those of exogenous wild type (FLAG-3xHA-ADA2B; Figure 6D). Interestingly, markedly lower protein levels of the exogenous mutated forms compared to the exogenous wild type were observed (Figure 6E). Exogenous expression of ADA2B also resulted in lower levels of the endogenous form, but this dominant negative effect was less prominent when the exogenous versions harbored the triple mutations.

**Figure 6.**
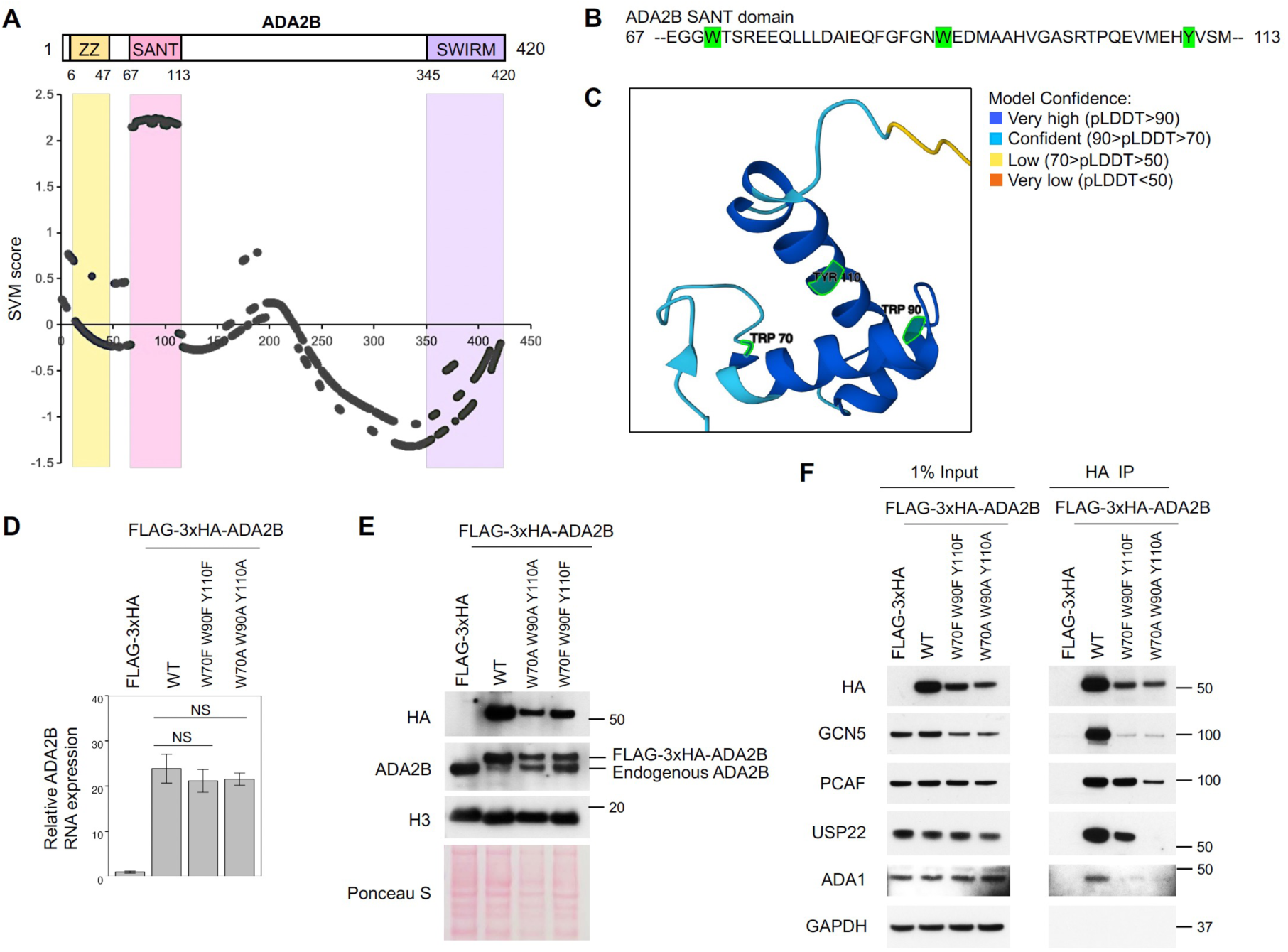
SANT domain of ADA2B is required for GCN5/PCAF interaction, SAGA incorporation, and ADA2B protein stability. **A)** Mapping of CRISPR KO hyper-sensitive regions in ADA2B protein by ProTiler (He et al. 2019a). Higher SVM scores indicate higher essentiality. Illustration of domains in ADA2B protein is shown at the top. **B)** Sequence of the SANT domain in ADA2B. The three predicted core hydrophobic residues (W70, W90, Y110) are highlighted in green. **C)** Structure of the SANT domain in ADA2B as predicted by AlphaFold (Jumper et al. 2021; Varadi et al. 2021). The three predicted core hydrophobic residues (W70, W90, Y110) are highlighted in green. AlphaFold generates a per-residue confidence score (pLDDT) ranging from 0 to 100. **D)** Relative RNA expression of ADA2B, including endogenous and exogenous levels, in MM.1S cell lines expressing FLAG-3xHA, FLAG-3xHA-ADA2B, FLAG-3xHA-ADA2B^W70A^ ^W90A^ ^Y110A^, or FLAG-3xHA-ADA2B^W70F^ ^W90F^ ^Y110F^ as determined by qRT-PCR. A one-way Analysis of Variance (ANOVA) was performed to analyze the differences among group means, followed by the Tukey HSD post hoc test to determine whether the mean difference between specific pairs of group are statistically significant. Not significant (NS) p > 0.05. **E)** Immunoblots showing levels of FLAG-3xHA-ADA2B and endogenous ADA2B in the same cell lines as in panel C. Histone H3 blot and Ponceau S staining are included as loading controls. **F)** Immunoblots showing co-IP results for FLAG-3xHA, FLAG-3xHA-ADA2B, FLAG-3xHA-ADA2B^W70A^ ^W90A^ ^Y110A^, and FLAG-3xHA-ADA2B^W70F^ ^W90F^ ^Y110F^ using the same cell lines as in panels C and D. GAPDH is included as a loading control in input samples.

No significant changes in MM cell viability were observed in cell lines expressing either FLAG-3xHA, FLAG-3xHA-ADA2B, FLAG-3xHA-ADA2B^W70F W90F Y110F^, or FLAG-3xHA-ADA2B^W70F W90F Y110F^ (Figure S7C). Since endogenous ADA2B is strongly inhibited in cells expressing FLAG-3xHA-ADA2B, the lack of changes in viability indicates that this epitope-tagged ADA2B (which was also used in our CUT&RUN) is fully functional. Indeed, as expected, CRISPR sgRNAs that simultaneously target both endogenous and exogenous ADA2B (sgADA2B) compromised cell viability, accompanied by decreased levels of GCN5, PCAF and MAF, as well as global H3K9ac (Figures S7D and S7E).

We hypothesized that overexpression of wild-type ADA2B displaced the endogenous protein from the KAT module, triggering turnover of the displaced protein. However, the triple mutations in the SANT domain likely weaken ADA2B–GCN5 interactions, such that mutant ADA2B is not stably incorporated into SAGA, rendering it less effective at displacing the endogenous protein. In agreement with this hypothesis, co-immunoprecipitations (co-IP) showed that FLAG-3xHA-ADA2B strongly associated with GCN5, but the two mutated versions did not (Figure 6F). Association with the SAGA core subunit ADA1 was also disrupted by the SANT domain mutations. Interestingly, in contrast to GCN5, PCAF associated with FLAG-3xHA-ADA2B^W70F W90F Y110F^ nearly as strongly as with FLAG-3xHA-ADA2B, whereas the alanine substitutions impaired interactions with both GCN5 and PCAF. The discrepancy in the effects of the mutations on interactions with GCN5 and PCAF suggests that interaction surfaces of ADA2B– GCN5 differs from those of ADA2B–PCAF. Association of USP22 with ADA2B was also compromised by the SANT domain triple mutations, especially with FLAG-3xHA-ADA2B^W70A W90A Y110A^ protein.

Collectively, these results demonstrate that the ADA2B SANT domain is required for SAGA integrity and KAT function. Loss of interactions with other SAGA components compromises ADA2B protein stability, presenting a vulnerability that could be targeted for therapeutic purposes.

## Discussion

Many histone modifying enzymes are part of multiple distinct complexes. Defining the division of labor between these complexes is imperative to understanding the proper regulation of gene expression programs both in normal cells and in disease states. Previous studies have elucidated important functions of GCN5 in various MYC-driven cancers (Chen et al. 2013; Yin et al. 2015; Majaz et al. 2016; Tzelepis et al. 2016; Farria et al. 2019; Mustachio et al. 2019; Farria et al. 2020; Oh et al. 2020). However, these studies did not address whether these functions were implemented through SAGA, ATAC, or both complexes. In this study, we demonstrate SAGA-specific ADA2B is required for MM cell growth, and repression of the SAGA KAT function through ADA2B depletion downregulates oncogenic transcription programs driven by MYC, E2F, and MAF. Our findings establish the critical role of the SAGA KAT module in the growth and survival of MM cells. However, we do not rule out that the ATAC complex may also have functions in MM.

The impact of ADA2B loss on MYC and MAF protein levels is post-transcriptional, as *MYC* and *MAF* RNA expression was not decreased. Additionally, inhibition of GCN5/PCAF KAT activity induced loss of these TFs, indicating acetylation has a direct or indirect impact on their stability. Acetylation of MYC by GCN5 and other KATs prevents ubiquitination and subsequent proteasomal degradation (Patel et al. 2004; Hurd et al. 2023). MAF protein stability and activity are known to be regulated by ubiquitination and phosphorylation (Chen et al. 2014; Herath et al. 2014; Xu et al. 2021), but acetylation of MAF has not yet been reported. Future work will ascertain whether MAF is a substrate for SAGA.

Our analyses indicate that MAF shares a majority of its gene targets with MYC. MYC exhibited a more robust binding to promoters of ADA2B-bound sites, whereas MAF predominantly bound to non-promoter regions, as observed by others (Katsarou et al. 2023). However, the extensive overlapping binding sites for ADA2B, MYC, and MAF at many common gene targets suggests these regulators function together in this context. Coregulation by MYC and MAF may serve as a redundant mechanism to enhance expression of these genes during MM development. Precedent for this cooperation exists in cholangiocarcinoma, where MYC and MAF interact directly with one another and also bind to E-boxes within their own promoters (Yang et al. 2016). Further studies are needed to determine whether these oncogenic TFs also physically interact in MM.

Integration of genes downregulated upon ADA2B loss and ADA2B binding sites identified cell cycle related genes and MTORC1 signaling genes among the ADA2B core targets. Deregulation of cell cycle progression is one of the key hallmarks of cancer and is consistent with MYC and MAF functions in MM (Chesi et al. 2008; Maes et al. 2017; Katsarou et al. 2023); MTORC1 signaling promotes tumor formation, proliferation, and metastasis (Li et al. 2014; Feng et al. 2020; Zou et al. 2020). Numerous cell cycle and MTOR pathway inhibitors are currently undergoing clinical and preclinical evaluation for MM therapy, but effectively targeting oncogenic TFs such as MYC and MAF remains a significant challenge. Targeting coactivators such as SAGA could offer an alternative therapeutic approach. Our studies show the SANT domain of ADA2B is critical for interaction with GCN5 as well as components of other SAGA modules. Development of small molecules or other strategies to compromise the functions of the SANT domain provides new avenues for therapy development.

Surprisingly, we found that substituting the hydrophobic core residues in the ADA2B SANT domain to phenylalanine did not disrupt interactions with PCAF, in contrast to its negative effect on interactions with GCN5. Correspondingly, we observed that mutations in the core residues led to decreased GCN5 protein levels likely due to its displacement from SAGA. Structural studies comparing GCN5– and PCAF-containing complexes are lacking, therefore the molecular basis for differences in the interaction structures of ADA2B with GCN5 as opposed to with PCAF is not yet clear. Our data represents one of the first few studies showing differential requirements of the KATs for associations with other SAGA complex subunits.

Of note, the SANT domain exists in other key transcriptional and chromatin regulators such as SMARCC1, SMRT, and EZH2 in humans (Boyer et al. 2002; Boyer et al. 2004; Weaver et al. 2019). EZH2 is an attractive drug target for various cancers including MM (Alzrigat et al. 2018a). EZH2 harbors two SANT domains, with SANT1 domain serving as a reader for unmodified histone H4. We also identified *EZH2* as a core gene target of ADA2B. These findings further support the potential of SANT domain inhibitors as possible new therapeutic approaches for MM and other cancers.

## Materials and Methods

### Cell lines

MM cell lines were purchased from American Type Culture Collection (ATCC) and cultured in RPMI-1640 supplemented with 10% fetal bovine serum (FBS), 100 U/ml penicillin and 100 µg/ml streptomycin (1% Pen-Strep). HEK-293T cells were provided by Dr. Han Xu lab (MD Anderson Cancer Center) and cultured in DMEM supplemented with 10% FBS and 1% Pen-Strep. Cell lines were tested regularly for mycoplasma (Lonza MycoAlert) to confirm negative infection.

### Transfection, viral transduction and inducible knockdown

Inducible hairpin-containing SMARTvector plasmids were obtained from Horizon Discovery/Dharmacon. Constitutive CRISPR sgRNA constructs were provided by Dr. Kimberly Stegmaier lab (Dana-Farber Cancer Institute). The following guide sequences were used: sgADA2B-4 GACAGGTGTGGTCTGTCACG, sgADA2B-5 GCTGGTAGCCGTGGTAGCGG, sgChr2-2 GGTGTGCGTATGAAGCAGTG, sgLacZ AACGGCGGATTGACCGTAAT.

Transfection was performed on HEK293T cells using the X-tremeGENE HP DNA Transfection Reagent (Roche) following manufacturer’s manual and viral supernatant was collected 48 hours after transfection. For spinoculation of MM cells, cell culture with viral particles and 8 µg/ml polybrene were centrifuged at 800g, 32^°^C for 1 hour. Transduced cells were selected by 1 µg/ml puromycin for 3 days. Expression of shRNA was induced by treatment of 1 µg/ml Dox to cell culture.

### Cell cycle analysis

Cells were washed in PBS, fixed in 70% ethanol, and incubated at –20^°^C overnight. Fixed cells were then centrifuged, washed in PBS, and resuspended in FxCycle PI/RNase staining solution (Invitrogen). After incubation at room temperature for 30 minutes, cells analyzed by flow cytometry using BD LSRFortessa. Results were analyzed by FlowJo software.

### Exogenous expression of ADA2B

Mutant versions of ADA2B were obtained by site-directed mutagenesis. FLAG and 3xHA tagged wild-type and mutated forms of ADA2B were cloned into pLVX-IRES-tdTomato vector, which was provided by Dr. Han Xu lab (MD Anderson Cancer Center). To generate MM cells expressing the tagged exogenous ADA2B, transfection and transduction were performed using the constructed plasmids. Red fluorescence (emitted by tdTomato) positive cells were sorted by flow cytometry using BD FACSAria Fusion.

### Immunoblotting and co-IP

For immunoblot analysis, cells were washed with PBS and subsequently lysed in RIPA buffer containing protease inhibitor cocktail (Roche). Protein samples were resolved by SDS–PAGE on 4-12% Nu-Page gels (Life Technologies) and transferred to PVDF membranes. The following antibodies were used: anti-ADA2B (Abcam, 1:1000 (discontinued) or Santa Cruz, 1:500), anti-GCN5L2 (Cell Signaling Technology, 1:1000), anti-PCAF (Cell Signaling Technology,1:1000), anti-ADA1 (Proteintech, 1:2000), anti-USP22 (Abcam, 1:2000), anti-MYC (Cell Signaling Technology, 1:1000), anti-MAF (Abcam, 1:1000), anti-E2F1 (Santa Cruz, 1:1000), anti-DP1 (Santa Cruz, 1:1000), anti-IRF4 (Santa Cruz, 1:500), anti-H3K9ac (Abcam, 1:1000), anti-H3 (Abcam, 1:20,000), anti-GAPDH (Millipore, 1:10,000), anti-β-Tubulin (Cell Signaling Technology, 1:1000), anti-HA (Cell Signaling Technology, 1:1000), and secondary horseradish peroxidase (HRP) conjugated antibodies (GE Healthcare). Amersham ECL Prime Western Blotting Detection Reagent (GE Healthcare) was used for chemiluminescent protein detection.

For co-IP, cells were lysed in IP lysis buffer (ThermoFisher) containing protease inhibitor cocktail (Roche) and phosphatase inhibitors (Roche). Equal amounts of protein lysates were used to incubate with pre-washed anti-HA magnetic beads (ThermoFisher) at 4^°^C for 3 hours, then washed 4 times and eluted with HA peptide (ThermoFisher) at 37^°^C for 10 minutes. Eluates and input samples were separated by SDS–PAGE and immunoblotted.

### Total RNA-seq and qRT-PCR

Total RNA was extracted from cell pellets using RNeasy Plus Mini Kit (Qiagen). The extracted RNA samples were treated with RNase-free DNase set (Qiagen). For RNA-seq, 1 μg of total RNA was used. Library preparation and sequencing were performed by the Advanced Technology Genomics Core at MD Anderson Cancer Center using Illumina NovaSeq6000 S1-200 flow cell and 100bp paired-end reading.

qRT-PCR was performed using the Power SYBR Green 1-step kit and the 7500 Fast Real-Time PCR System (Applied Biosystems). Relative expression was calculated using the comparative Ct method with *Actin B* (*ACTB*) as an internal control. The following primer sequences were used: ADA2B-F GCTACCACGGCTACCAGC, ADA2B-R AGCCATATCTTCCCAGTTTCCG, ADA2B-F2 ATGTACGTGCGGAAGCTGAA, ADA2B-R2 AGCTCCTTCTCCTCCTTGGT, ACTB-F AGAGCTACGAGCTGCCTGAC, ACTB-R AGCACTGTGTTGGCGTACAG.

### ATAC-seq

The Tn5 transposition reaction was performed using the Illumina Tagment DNA TDE1 Enzyme and Buffer Kit (Illumina). For each sample 100,000 cells were collected and washed with cold PBS. A resuspension buffer (10 nM Tris-HCl, pH7.5, 10 mM NaCl, 3 mM MgCl_2_) was used for subsequent steps. Cells were lysed in 100 μl lysis buffer (0.1% NP-40, 0.1% Tween-20, and 0.01% Digitonin in resuspension buffer) on ice for 3 minutes, then washed with 1 ml wash buffer (0.1% Tween-20 in resuspension buffer).

Lysates were cleared by centrifugation at 500g, 4^°^C for 10 minutes. The nuclear pellet was resuspended in 100 μl transposition reaction mix (50 μl Tagment DNA Buffer, 33 μl PBS, 1 μl 10% Tween-20, 1 μl 1% Digitonin, 5 μl TDE1, 10 μl nuclease-free water) and incubated at 37^°^C on a thermomixer with 1,000 rpm for 30 minutes. DNA was isolated using the Qiagen MinElute Reaction Cleanup Kit (Qiagen). Library was prepared with NEBNext High-Fidelity 2X PCR Master Mix (NEB) and SYBR Green I (ThermoFisher), then QC analyzed by the Epigenomics Profiling Core at MD Anderson Cancer Center. Sequencing was performed at the Advanced Technology Genomics Core at MD Anderson Cancer Center using Illumina NovaSeq6000 S1-100 flow cell and 50bp paired-end reading.

### CUT&RUN

MM.1S cells expressing either FLAG-3xHA or FLAG-3xHA-ADA2B were subjected to light crosslinking with 0.1% formaldehyde for 1 minute, followed by quenching with 125 mM glycine. Remaining steps of CUT&RUN were performed by the Epigenomics Profiling Core at MD Anderson Cancer Center according to published protocols (Meers et al. 2019) with some modifications. Briefly, 500,000 cells were immobilized to activated Concanavalin A-coated magnetic beads (Bangs Laboratories) followed by permeabilization with wash buffer containing Digitonin (Promega). Samples were incubated with rabbit IgG (Millipore), H3K9ac (Diagenode) and HA (Cell Signaling Technology) antibodies overnight at 4°C. Targeted chromatin digestion was achieved by pAG-MNase (EpiCypher) binding for 10 minutes at room temperature followed by incubation with CaCl_2_ at 4°C. DNA fragments were purified using MinElute columns (Qiagen). Libraries were prepared using NEBNext Ultra II DNA Library prep kit (New England Biolabs) following manufacturer’s instructions for H3K9ac CUT&RUN DNA, and a modified library preparation protocol for HA CUT&RUN DNA as described previously (Liu et al. 2018). Libraries were sequenced at the Advanced Technology Genomics Core at MD Anderson Cancer Center using Illumina NovaSeq 6000 SP-100 flow cell to obtain 50bp paired-end reads.

## Data availability

All new genome-scale omics datasets generated in this study are made publicly available at the Gene Expression Omnibus (GEO) repository https://www.ncbi.nlm.nih.gov/geo/ as GSE262885, GSE262886, and GSE262887. The following secure tokens have been created to allow review of records while they remain in private status: ajehamsixluzpmv for GSE262885, adefwkcidrolfgz for GSE262886, and enijycugpbexhul for GSE262887. These datasets will become public upon publication.

## Acknowledgments

We thank Dr. Abhinav Jain and the Epigenomics Profiling Core at MD Anderson Cancer Center for technical assistance. We also thank Dr. Kimberly Stegmaier lab for gifting the CRISPR sgRNA constructs, Dr. Han Xu lab for gifting the pLVX-IRES-tdTomato vector, and Dr. Wei He for assistance with the ProTiler software. We are grateful to Dr. Gheath Al-Atrash and Mao Zhang for discussions regarding MM pathology. We thank members of the Dent lab for critical discussions regarding this work. Y-JCC was supported by the Scholar Fellowship Award from the Center for Cancer Epigenetics at The University of Texas MD Anderson Cancer Center. This study was supported by NIH grants R35 GM131678 awarded to SYRD and 1S10OD024977-01 awarded to the Advanced Technology Genomics Core. The Advanced Technology Genomics Core was supported in part by The University of Texas MD Anderson Cancer Center and P30CA016672.

## Author Contributions

Y-JCC and SYRD conceived and directed the project. Y-JCC designed and performed the experiments, analyzed data and interpreted results. BGB performed gene expression and protein studies including qRT-PCR and co-IP, and helped with data analysis and interpretation. YL performed bioinformatics analyses of the sequencing data with assistance from KL. Y-JCC wrote the original draft of the manuscript with SYRD. All authors reviewed and approved the final version of the manuscript. SYRD acquired the funding.

## References

1. Alvarez-Benayas J, Trasanidis N, Katsarou A, Ponnusamy K, Chaidos A, May PC, Xiao X, Bua M, Atta M, Roberts IAG et al. 2021. Chromatin-based, in cis and in trans regulatory rewiring underpins distinct oncogenic transcriptomes in multiple myeloma. Nat Commun 12: 5450.

2. Alzrigat M, Jernberg-Wiklund H, Licht JD. 2018a. Targeting EZH2 in Multiple Myeloma-Multifaceted Anti-Tumor Activity. Epigenomes 2.

3. Alzrigat M, Parraga AA, Jernberg-Wiklund H. 2018b. Epigenetics in multiple myeloma: From mechanisms to therapy. Semin Cancer Biol 51: 101–115.

4. Arede L, Foerner E, Wind S, Kulkarni R, Domingues AF, Giotopoulos G, Kleinwaechter S, Mollenhauer-Starkl M, Davison H, Chandru A et al. 2022. KAT2A complexes ATAC and SAGA play unique roles in cell maintenance and identity in hematopoiesis and leukemia. Blood Adv 6: 165–180.

5. Bergsagel PL, Kuehl WM. 2003. Critical roles for immunoglobulin translocations and cyclin D dysregulation in multiple myeloma. Immunological Reviews 194: 96–104.

6. Boyer LA, Langer MR, Crowley KA, Tan S, Denu JM, Peterson CL. 2002. Essential Role for the SANT Domain in the Functioning of Multiple Chromatin Remodeling Enzymes. Molecular Cell 10: 935–942.

7. Boyer LA, Latek RR, Peterson CL. 2004. The SANT domain: a unique histone-tail-binding module? Nature Reviews Molecular Cell Biology 5: 158–163.

8. Brownell JE, Allis CD. 1995. An activity gel assay detects a single, catalytically active histone acetyltransferase subunit in Tetrahymena macronuclei. Proc Natl Acad Sci U S A 92: 6364–6368.

9. Brownell JE, Zhou J, Ranalli T, Kobayashi R, Edmondson DG, Roth SY, Allis CD. 1996. Tetrahymena histone acetyltransferase A: a homolog to yeast Gcn5p linking histone acetylation to gene activation. Cell 84: 843–851.

10. Chen G, Xu X, Tong J, Han K, Zhang Z, Tang J, Li S, Yang C, Li J, Cao B et al. 2014. Ubiquitination of the transcription factor c-MAF is mediated by multiple lysine residues. Int J Biochem Cell Biol 57: 157–166.

11. Chen L, Wei T, Si X, Wang Q, Li Y, Leng Y, Deng A, Chen J, Wang G, Zhu S et al. 2013. Lysine acetyltransferase GCN5 potentiates the growth of non-small cell lung cancer via promotion of E2F1, cyclin D1, and cyclin E1 expression. J Biol Chem 288: 14510–14521.

12. Chen Y-JC, Dent SYR. 2021. Conservation and diversity of the eukaryotic SAGA coactivator complex across kingdoms. Epigenetics & Chromatin 14: 26.

13. Chen YC, Koutelou E, Dent SYR. 2022. Now open: Evolving insights to the roles of lysine acetylation in chromatin organization and function. Mol Cell 82: 716–727.

14. Chesi M, Robbiani DF, Sebag M, Chng WJ, Affer M, Tiedemann R, Valdez R, Palmer SE, Haas SS, Stewart AK et al. 2008. AID-dependent activation of a MYC transgene induces multiple myeloma in a conditional mouse model of post-germinal center malignancies. Cancer Cell 13: 167–180.

15. Downey M. 2021. Non-histone protein acetylation by the evolutionarily conserved GCN5 and PCAF acetyltransferases. Biochim Biophys Acta Gene Regul Mech 1864: 194608.

16. Farria AT, Mustachio LM, Akdemir ZHC, Dent SYR. 2019. GCN5 HAT inhibition reduces human Burkitt lymphoma cell survival through reduction of MYC target gene expression and impeding BCR signaling pathways. Oncotarget 10: 5847–5858.

17. Farria AT, Plummer JB, Salinger AP, Shen J, Lin K, Lu Y, McBride KM, Koutelou E, Dent SYR. 2020. Transcriptional Activation of MYC-Induced Genes by GCN5 Promotes B-cell Lymphomagenesis. Cancer Res 80: 5543–5553.

18. Feng G, Arima Y, Midorikawa K, Kobayashi H, Oikawa S, Zhao W, Zhang Z, Takeuchi K, Murata M. 2023. Knockdown of TFRC suppressed the progression of nasopharyngeal carcinoma by downregulating the PI3K/Akt/mTOR pathway. Cancer Cell Int 23: 185.

19. Feng Y, Chen X, Cassady K, Zou Z, Yang S, Wang Z, Zhang X. 2020. The Role of mTOR Inhibitors in Hematologic Disease: From Bench to Bedside. Front Oncol 10: 611690.

20. Fulciniti M, Lin CY, Samur MK, Lopez MA, Singh I, Lawlor MA, Szalat RE, Ott CJ, Avet-Loiseau H, Anderson KC et al. 2018. Non-overlapping Control of Transcriptome by Promoter– and Super-Enhancer-Associated Dependencies in Multiple Myeloma. Cell Rep 25: 3693–3705 e3696.

21. Gamper AM, Kim J, Roeder RG. 2009. The STAGA subunit ADA2b is an important regulator of human GCN5 catalysis. Mol Cell Biol 29: 266–280.

22. Grant PA, Duggan L, Côté J, Roberts SM, Brownell JE, Candau R, Ohba R, Owen-Hughes T, Allis CD, Winston F et al. 1997. Yeast Gcn5 functions in two multisubunit complexes to acetylate nucleosomal histones: characterization of an Ada complex and the SAGA (Spt/Ada) complex. Genes Dev 11: 1640–1650.

23. Guelman S, Kozuka K, Mao Y, Pham V, Solloway MJ, Wang J, Wu J, Lill JR, Zha J. 2009. The double-histone-acetyltransferase complex ATAC is essential for mammalian development. Mol Cell Biol 29: 1176–1188.

24. Hanamura I, Huang Y, Zhan F, Barlogie B, Shaughnessy J. 2006. Prognostic value of Cyclin D2 mRNA expression in newly diagnosed multiple myeloma treated with high-dose chemotherapy and tandem autologous stem cell transplantations. Leukemia 20: 1288–1290.

25. He W, Zhang L, Villarreal OD, Fu R, Bedford E, Dou J, Patel AY, Bedford MT, Shi X, Chen T et al. 2019a. De novo identification of essential protein domains from CRISPR-Cas9 tiling-sgRNA knockout screens. Nature Communications 10: 4541.

26. He Y, Luo Y, Zhang D, Wang X, Zhang P, Li H, Ejaz S, Liang S. 2019b. PGK1-mediated cancer progression and drug resistance. Am J Cancer Res 9: 2280–2302.

27. Herath NI, Rocques N, Garancher A, Eychene A, Pouponnot C. 2014. GSK3-mediated MAF phosphorylation in multiple myeloma as a potential therapeutic target. Blood Cancer J 4: e175.

28. Herbst DA, Esbin MN, Louder RK, Dugast-Darzacq C, Dailey GM, Fang Q, Darzacq X, Tjian R, Nogales E. 2021. Structure of the human SAGA coactivator complex. Nat Struct Mol Biol 28: 989–996.

29. Hernández Borrero LJ, El-Deiry WS. 2021. Tumor suppressor p53: Biology, signaling pathways, and therapeutic targeting. Biochimica et Biophysica Acta (BBA) – Reviews on Cancer 1876: 188556.

30. Hirsch CL, Coban Akdemir Z, Wang L, Jayakumaran G, Trcka D, Weiss A, Hernandez JJ, Pan Q, Han H, Xu X et al. 2015. Myc and SAGA rewire an alternative splicing network during early somatic cell reprogramming. Genes Dev 29: 803–816.

31. Hurd M, Pino J, Jang K, Allevato MM, Vorontchikhina M, Ichikawa W, Zhao Y, Gates R, Villalpando E, Hamilton MJ et al. 2023. MYC acetylated lysine residues drive oncogenic cell transformation and regulate select genetic programs for cell adhesion-independent growth and survival. Genes Dev 37: 865–882.

32. Hurt EM, Wiestner A, Rosenwald A, Shaffer AL, Campo E, Grogan T, Bergsagel PL, Kuehl WM, Staudt LM. 2004. Overexpression of c-maf is a frequent oncogenic event in multiple myeloma that promotes proliferation and pathological interactions with bone marrow stroma. Cancer Cell 5: 191–199.

33. Ianari A, Gallo R, Palma M, Alesse E, Gulino A. 2004. Specific role for p300/CREB-binding protein-associated factor activity in E2F1 stabilization in response to DNA damage. J Biol Chem 279: 30830–30835.

34. Jumper J, Evans R, Pritzel A, Green T, Figurnov M, Ronneberger O, Tunyasuvunakool K, Bates R, Žídek A, Potapenko A, et al. 2021. Highly accurate protein structure prediction with AlphaFold. Nature 596: 583–589.

35. Katsarou A, Trasanidis N, Ponnusamy K, Kostopoulos IV, Alvarez-Benayas J, Papaleonidopoulou F, Keren K, Sabbattini PMR, Feldhahn N, Papaioannou M et al. 2023. MAF functions as a pioneer transcription factor that initiates and sustains myelomagenesis. Blood Adv 7: 6395–6410.

36. Kiss Z, Mudryj M, Ghosh PM. 2020. Non-circadian aspects of BHLHE40 cellular function in cancer. Genes Cancer 11: 1–19.

37. Knoepfler PS, Zhang Xy, Cheng PF, Gafken PR, McMahon SB, Eisenman RN. 2006. Myc influences global chromatin structure. The EMBO Journal 25: 2723–2734.

38. Kusch T, Guelman S, Abmayr SM, Workman JL. 2003. Two Drosophila Ada2 homologues function in different multiprotein complexes. Mol Cell Biol 23: 3305–3319.

39. Li J, Zhu J, Cao B, Mao X. 2014. The mTOR signaling pathway is an emerging therapeutic target in multiple myeloma. Curr Pharm Des 20: 125–135.

40. Lin CY, Lovén J, Rahl PB, Paranal RM, Burge CB, Bradner JE, Lee TI, Young RA. 2012. Transcriptional amplification in tumor cells with elevated c-Myc. Cell 151: 56–67.

41. Liu N, Hargreaves VV, Zhu Q, Kurland JV, Hong J, Kim W, Sher F, Macias-Trevino C, Rogers JM, Kurita R et al. 2018. Direct Promoter Repression by BCL11A Controls the Fetal to Adult Hemoglobin Switch. Cell 173: 430–442.e417.

42. Maes A, Menu E, Veirman K, Maes K, Vand Erkerken K, De Bruyne E. 2017. The therapeutic potential of cell cycle targeting in multiple myeloma. Oncotarget 8: 90501–90520.

43. Majaz S, Tong Z, Peng K, Wang W, Ren W, Li M, Liu K, Mo P, Li W, Yu C. 2016. Histone acetyl transferase GCN5 promotes human hepatocellular carcinoma progression by enhancing AIB1 expression. Cell Biosci 6: 47.

44. Meers MP, Bryson TD, Henikoff JG, Henikoff S. 2019. Improved CUT&RUN chromatin profiling tools. eLife 8: e46314.

45. Mo Z, Zhang S, Zhang S. 2020. A Novel Signature Based on mTORC1 Pathway in Hepatocellular Carcinoma. Journal of Oncology 2020: 8291036.

46. Motohashi H, Shavit JA, Igarashi K, Yamamoto M, Engel JD. 1997. The world according to Maf. Nucleic Acids Res 25: 2953–2959.

47. Muratoglu S, Georgieva S, Pápai G, Scheer E, Enünlü I, Komonyi O, Cserpán I, Lebedeva L, Nabirochkina E, Udvardy A et al. 2003. Two different Drosophila ADA2 homologues are present in distinct GCN5 histone acetyltransferase-containing complexes. Mol Cell Biol 23: 306–321.

48. Mustachio LM, Roszik J, Farria AT, Guerra K, Dent SY. 2019. Repression of GCN5 expression or activity attenuates c-MYC expression in non-small cell lung cancer. Am J Cancer Res 9: 1830–1845.

49. Muylaert C, Van Hemelrijck LA, Maes A, De Veirman K, Menu E, Vanderkerken K, De Bruyne E. 2022. Aberrant DNA methylation in multiple myeloma: A major obstacle or an opportunity? Front Oncol 12: 979569.

50. Neri P, Ren L, Azab AK, Brentnall M, Gratton K, Klimowicz AC, Lin C, Duggan P, Tassone P, Mansoor A et al. 2011. Integrin β7-mediated regulation of multiple myeloma cell adhesion, migration, and invasion. Blood 117: 6202–6213.

51. Oh JH, Lee J-Y, Kim KH, Kim CY, Jeong DS, Cho Y, Nam KT, Kim MH. 2020. Elevated GCN5 expression confers tamoxifen resistance by upregulating AIB1 expression in ER-positive breast cancer. Cancer Letters 495: 145–155.

52. Pacini C, Dempster JM, Boyle I, Gonçalves E, Najgebauer H, Karakoc E, van der Meer D, Barthorpe A, Lightfoot H, Jaaks P et al. 2021. Integrated cross-study datasets of genetic dependencies in cancer. Nat Commun 12: 1661.

53. Patel JH, Du Y, Ard PG, Phillips C, Carella B, Chen CJ, Rakowski C, Chatterjee C, Lieberman PM, Lane WS et al. 2004. The c-MYC oncoprotein is a substrate of the acetyltransferases hGCN5/PCAF and TIP60. Mol Cell Biol 24: 10826–10834.

54. Qian X, Yang Y, Deng Y, Liu Y, Zhou Y, Han F, Xu Y, Yuan H. 2023. SETDB1 induces lenalidomide resistance in multiple myeloma cells via epithelial–mesenchymal transition and PI3K/AKT pathway activation. Exp Ther Med 25: 274.

55. Riss A, Scheer E, Joint M, Trowitzsch S, Berger I, Tora L. 2015. Subunits of ADA-two-A-containing (ATAC) or Spt-Ada-Gcn5-acetyltrasferase (SAGA) Coactivator Complexes Enhance the Acetyltransferase Activity of GCN5. J Biol Chem 290: 28997–29009.

56. Sarin V, Yu K, Ferguson ID, Gugliemini O, Nix MA, Hann B, Sirota M, Wiita AP. 2020. Evaluating the efficacy of multiple myeloma cell lines as models for patient tumors via transcriptomic correlation analysis. Leukemia 34: 2754–2765.

57. Shan J, Wang Z, Mo Q, Long J, Fan Y, Cheng L, Zhang T, Liu X, Wang X. 2022. Ribonucleotide reductase M2 subunit silencing suppresses tumorigenesis in pancreatic cancer via inactivation of PI3K/AKT/mTOR pathway. Pancreatology 22: 401–413.

58. Suganuma T, Gutiérrez JL, Li B, Florens L, Swanson SK, Washburn MP, Abmayr SM, Workman JL. 2008. ATAC is a double histone acetyltransferase complex that stimulates nucleosome sliding. Nat Struct Mol Biol 15: 364–372.

59. Sun J, Paduch M, Kim SA, Kramer RM, Barrios AF, Lu V, Luke J, Usatyuk S, Kossiakoff AA, Tan S. 2018. Structural basis for activation of SAGA histone acetyltransferase Gcn5 by partner subunit Ada2. Proc Natl Acad Sci U S A 115: 10010–10015.

60. Trisciuoglio D, Ragazzoni Y, Pelosi A, Desideri M, Carradori S, Gabellini C, Maresca G, Nescatelli R, Secci D, Bolasco A et al. 2012. CPTH6, a thiazole derivative, induces histone hypoacetylation and apoptosis in human leukemia cells. Clin Cancer Res 18: 475–486.

61. Tzelepis K, Koike-Yusa H, De Braekeleer E, Li Y, Metzakopian E, Dovey OM, Mupo A, Grinkevich V, Li M, Mazan M et al. 2016. A CRISPR Dropout Screen Identifies Genetic Vulnerabilities and Therapeutic Targets in Acute Myeloid Leukemia. Cell Rep 17: 1193–1205.

62. Varadi M, Anyango S, Deshpande M, Nair S, Natassia C, Yordanova G, Yuan D, Stroe O, Wood G, Laydon A et al. 2021. AlphaFold Protein Structure Database: massively expanding the structural coverage of protein-sequence space with high-accuracy models. Nucleic Acids Research 50: D439–D444.

63. Wang YL, Faiola F, Xu M, Pan S, Martinez E. 2008. Human ATAC Is a GCN5/PCAF-containing acetylase complex with a novel NC2-like histone fold module that interacts with the TATA-binding protein. J Biol Chem 283: 33808–33815.

64. Weaver TM, Liu J, Connelly KE, Coble C, Varzavand K, Dykhuizen EC, Musselman CA. 2019. The EZH2 SANT1 domain is a histone reader providing sensitivity to the modification state of the H4 tail. Scientific Reports 9: 987.

65. Wu D, Qiu Y, Jiao Y, Qiu Z, Liu D. 2020. Small Molecules Targeting HATs, HDACs, and BRDs in Cancer Therapy. Front Oncol 10: 560487.

66. Xu W, Edmondson DG, Evrard YA, Wakamiya M, Behringer RR, Roth SY. 2000. Loss of Gcn5l2 leads to increased apoptosis and mesodermal defects during mouse development. Nat Genet 26: 229–232.

67. Xu Y, Xu M, Tong J, Tang X, Chen J, Chen X, Zhang Z, Cao B, Stewart AK, Moran MF et al. 2021. Targeting the Otub1/c-Maf axis for the treatment of multiple myeloma. Blood 137: 1478–1490.

68. Yamauchi T, Yamauchi J, Kuwata T, Tamura T, Yamashita T, Bae N, Westphal H, Ozato K, Nakatani Y. 2000. Distinct but overlapping roles of histone acetylase PCAF and of the closely related PCAF-B/GCN5 in mouse embryogenesis. Proc Natl Acad Sci U S A 97: 11303–11306.

69. Yang H, Liu T, Wang J, Li TW, Fan W, Peng H, Krishnan A, Gores GJ, Mato JM, Lu SC. 2016. Deregulated methionine adenosyltransferase α1, c-Myc, and Maf proteins together promote cholangiocarcinoma growth in mice and humans(‡). Hepatology 64: 439–455.

70. Yin YW, Jin HJ, Zhao W, Gao B, Fang J, Wei J, Zhang DD, Zhang J, Fang D. 2015. The Histone Acetyltransferase GCN5 Expression Is Elevated and Regulated by c-Myc and E2F1 Transcription Factors in Human Colon Cancer. Gene Expr 16: 187–196.

71. Zeller KI, Zhao X, Lee CW, Chiu KP, Yao F, Yustein JT, Ooi HS, Orlov YL, Shahab A, Yong HC et al. 2006. Global mapping of c-Myc binding sites and target gene networks in human B cells. Proc Natl Acad Sci U S A 103: 17834–17839.

72. Zhan F, Barlogie B, Arzoumanian V, Huang Y, Williams DR, Hollmig K, Pineda-Roman M, Tricot G, van Rhee F, Zangari M et al. 2007. Gene-expression signature of benign monoclonal gammopathy evident in multiple myeloma is linked to good prognosis. Blood 109: 1692–1700.

73. Zhan F, Huang Y, Colla S, Stewart JP, Hanamura I, Gupta S, Epstein J, Yaccoby S, Sawyer J, Burington B et al. 2006. The molecular classification of multiple myeloma. Blood 108: 2020–2028.

74. Zhao R, Tian L, Zhao B, Sun Y, Cao J, Chen K, Li F, Li M, Shang D, Liu M. 2020. FADS1 promotes the progression of laryngeal squamous cell carcinoma through activating AKT/mTOR signaling. Cell Death & Disease 11: 272.

75. Zou Z, Tao T, Li H, Zhu X. 2020. mTOR signaling pathway and mTOR inhibitors in cancer: progress and challenges. Cell & Bioscience 10: 31.

