## Supplemental Material for "The SAGA acetyltransferase module is required for the maintenance of MAF and MYC oncogenic gene expression programs in multiple myeloma"

**Supplemental Material includes:**

**Supplemental Material and Methods**

**Supplemental References**

**Supplemental Figures S1-S7**

**Supplemental Table S1:** Lists of downregulated genes and genes with decreased accessibility at both 3 days and 5 days of ADA2B knockdown.

**Supplemental Table S2:** Lists of ADA2B core gene targets at 3 days and 5 days of ADA2B knockdown.

### Supplemental Material and Methods

#### RNA-seq data analysis

***Mapping***: Adapter sequences were removed from the 3' ends of reads using Trim Galore! (version 0.6.5) and cutadapt (version 2.8) (Martin 2011). The reads were then aligned to the human genome (hg38) using TopHat (version 2.0.10) (Kim et al. 2013). ***Differential expression***: The number of fragments for each known gene from GENCODE (Frankish et al. 2021) Release 37 was quantified using htseq-count from the HTSeq package (version 0.11.0) (Anders et al. 2015). Genes with fewer than 10 fragments in all samples were excluded before the differential expression analysis. The statistical assessment of differential expression between conditions was performed using the R/Bioconductor package edgeR (version 3.8.6) (Robinson et al. 2010). ***Gene set enrichment analysis***: Gene set enrichment analysis was conducted using the GSEA software (Subramanian et al. 2005) against the Molecular Signatures Database (MSigDB) collections H and C2 gene sets (Castanza et al. 2023). The genes were ranked from the most upregulated to the most downregulated based on $-\log_{10}$(pvalue), with positive and negative signs representing upregulation or downregulation, respectively. Subsequently, GSEA was performed using the pre-ranked gene list, with all other parameters left as default.

#### ATAC-seq data analysis

***Mapping***: Adapter sequences were removed from the 3' ends of reads using Trim Galore! (version 0.6.5) and cutadapt (version 2.8). The reads were then mapped to the human genome (hg38) using Bowtie (version 1.1.2) (Langmead et al. 2009) with the following parameters: "--allow-contain --maxins 2000 -v 2 -m 1 --best --strata". To avoid PCR bias, only one copy of multiple fragments mapped to the same genomic position was retained for further analysis. After the removal of fragments from chrM, for each fragment, the 5' end was offset by +4 bp and the 3' end was offset by -5 bp to adjust both ends to represent the center of a transposon binding event. ***Peak calling***: The replicates of each condition were merged to represent the merged sample of that condition. For each merged and individual sample, peaks were identified using MACS2 (version 2.2.7.1) (Zhang et al. 2008) without using any control. Each binding event (i.e., the 5' or 3' end of a fragment) was smoothed by 73 bp (i.e., extended 36 bp upstream and 36 bp downstream from the event center). MACS2 was configured to call peaks from the pile-up of smoothed binding events, and a q-value cutoff of 0.001 was applied. The peaks that overlapped with ENCODE blacklisted regions (Amemiya et al. 2019) were removed. The peaks identified in the merged sample and overlapping with peaks in both replicates were considered representative of that condition and used for differential analysis. ***Differential peak analysis***: peaks for all the conditions were merged, and the number of transposon binding events within these merged peaks was counted for each sample. Merged peaks with less than 10 binding events in all samples were removed. The resulting count table was used to identify differential peaks using the R/Bioconductor package edgeR (Robinson et al. 2010). The numbers of binding events within the common peaks of all the conditions were used as the library sizes in edgeR. Peaks with a false discovery rate (FDR) ≤ 0.05 and a fold change (FC) ≥ 1.5 were identified as differential peaks. ***Signal track***: each transposon binding event was smoothed to a length of 73 bp, spanning from -36 bp to +36 bp around the center. The count of binding events covering each genomic position was multiplied by $1x10^7$ divided by the library size used in edgeR. These values were then averaged over a 10 bp resolution. The resulting averaged values were displayed using the Integrative Genomics Viewer (IGV) (Thorvaldsdóttir et al. 2013).

**CUT&RUN data analysis**

***Mapping***: Adapter sequences were removed from the 3' ends of reads using Trim Galore! (version 0.6.5) and cutadapt (version 2.8). The reads were then mapped to the human genome (hg38) using Bowtie (version 1.1.2) (Castanza et al. 2023) with the following parameters: ""--allow-contain --maxins 2000 -v 2 -m 1 --best --strata". To avoid PCR bias, only one copy of multiple fragments mapped to the same genomic position was retained for further analysis. Moreover, fragments longer than 800 bp (less than 1% of the total) were excluded from subsequent analysis. ***Peak calling for ADA2B***: peaks were identified using MACS2 (version 2.2.7.1) by comparing against the corresponding IgG sample. MACS2 was configured to call peaks from the pile-up of 151 bp around the middle point of each fragment. Since the two replicates of ADA2B exhibited different signal qualities, the final set of ADA2B peaks was determined by selecting the peaks that satisfied the following criteria: they were called in the ADA2B replicate with good signal at q-value 0.05, overlapped with the peaks called in the compromised replicate at p-value 0.01, and did not overlap with ENCODE blacklisted regions. ***Signal landscape***: To generate the signal landscape for CUT&RUN, each fragment was resized to a length of 151 bp, spanning from -75 bp to +75 bp around its midpoint. The count of fragments covering each genomic position was multiplied by $1x10^{-7}$ divided by the total number of mapped fragments. These values were then averaged over a 10 bp resolution. The resulting averaged values were displayed using IGV. ***Heatmap and metaplot***: Heatmaps were generated around the summits of ADA2B peaks or the centers of merged peaks from ADA2B, MAF, and MYC. The region spanning 10 kb upstream to 10 kb downstream from the summit or center of each peak was divided into 100 bp bins. For each sample, the FPKM or RPKM (fragments/reads per kilobase per million fragments/reads) values were calculated for all the bins and subtracted by the FPKM or RPKM values of the corresponding IgG or input sample. For CUT&RUN and ChIP-Seq samples, normalization was conducted per million total mapped fragments/reads. For ATAC-Seq samples, normalization was conducted per million fragments used in edgeR for differential analysis. The values were then averaged over all replicates, if available. Subsequently, for each data type and condition, all values were divided by the root mean square (rms) value to make value scales of all conditions similar. For the four ATAC-Seq conditions, their values were scaled together to preserve the original signal intensity contrast. The final value table for each data type and condition was visualized in a heatmap by the R function heatmap.2. For the metaplot, the resulting value table for each data type and condition was averaged over peaks within each group and plotted accordingly.

**Processing of public ChIP-seq data**

The raw fastq files of MAF and MYC ChIP-Seq data were downloaded from the Sequence Read Archive (SRA) with the following accession numbers: SRR20993552 (MAF), SRR20993551 (input for MAF), SRR444479 (MYC), and SRR444481 (input for MYC). A similar mapping and peak calling procedure as that for CUT&RUN was employed. Briefly, the reads were mapped to the human genome (hg38) using Bowtie with the following parameters: "-v 2 -m 1 --best --strata". To avoid PCR bias, only one copy of multiple fragments mapped to the same genomic position was retained. Peaks were identified using MACS2 by comparing against the corresponding input sample with a q-value cutoff of 0.05. The peaks that overlapped with ENCODE blacklisted regions were removed.

**Supplemental References**


Amemiya HM, Kundaje A, Boyle AP. 2019. The ENCODE Blacklist: Identification of Problematic Regions of the Genome. *Sci Rep* **9**: 9354.

Anders S, Pyl PT, Huber W. 2015. HTSeq--a Python framework to work with high-throughput sequencing data. *Bioinformatics* **31**: 166-169.

Castanza AS, Recla JM, Eby D, Thorvaldsdóttir H, Bult CJ, Mesirov JP. 2023. Extending support for mouse data in the Molecular Signatures Database (MSigDB). *Nat Methods* **20**: 1619-1620.

Frankish A, Diekhans M, Jungreis I, Lagarde J, Loveland JE, Mudge JM, Sisu C, Wright JC, Armstrong J, Barnes I et al. 2021. GENCODE 2021. *Nucleic Acids Res* **49**: D916-d923.

Kim D, Pertea G, Trapnell C, Pimentel H, Kelley R, Salzberg SL. 2013. TopHat2: accurate alignment of transcriptomes in the presence of insertions, deletions and gene fusions. *Genome Biol* **14**: R36.

Langmead B, Trapnell C, Pop M, Salzberg SL. 2009. Ultrafast and memory-efficient alignment of short DNA sequences to the human genome. *Genome Biol* **10**: R25.

Martin M. 2011. Cutadapt removes adapter sequences from high-throughput sequencing reads. *2011* **17**: 3.

Robinson MD, McCarthy DJ, Smyth GK. 2010. edgeR: a Bioconductor package for differential expression analysis of digital gene expression data. *Bioinformatics* **26**: 139-140.

Subramanian A, Tamayo P, Mootha VK, Mukherjee S, Ebert BL, Gillette MA, Paulovich A, Pomeroy SL, Golub TR, Lander ES et al. 2005. Gene set enrichment analysis:

a knowledge-based approach for interpreting genome-wide expression profiles. *Proc Natl Acad Sci U S A* **102**: 15545-15550.

Thorvaldsdóttir H, Robinson JT, Mesirov JP. 2013. Integrative Genomics Viewer (IGV): high-performance genomics data visualization and exploration. *Brief Bioinform* **14**: 178-192.

Zhang Y, Liu T, Meyer CA, Eeckhoute J, Johnson DS, Bernstein BE, Nusbaum C, Myers RM, Brown M, Li W et al. 2008. Model-based analysis of ChIP-Seq (MACS). *Genome Biol* **9**: R137.

Supplemental Figure S1

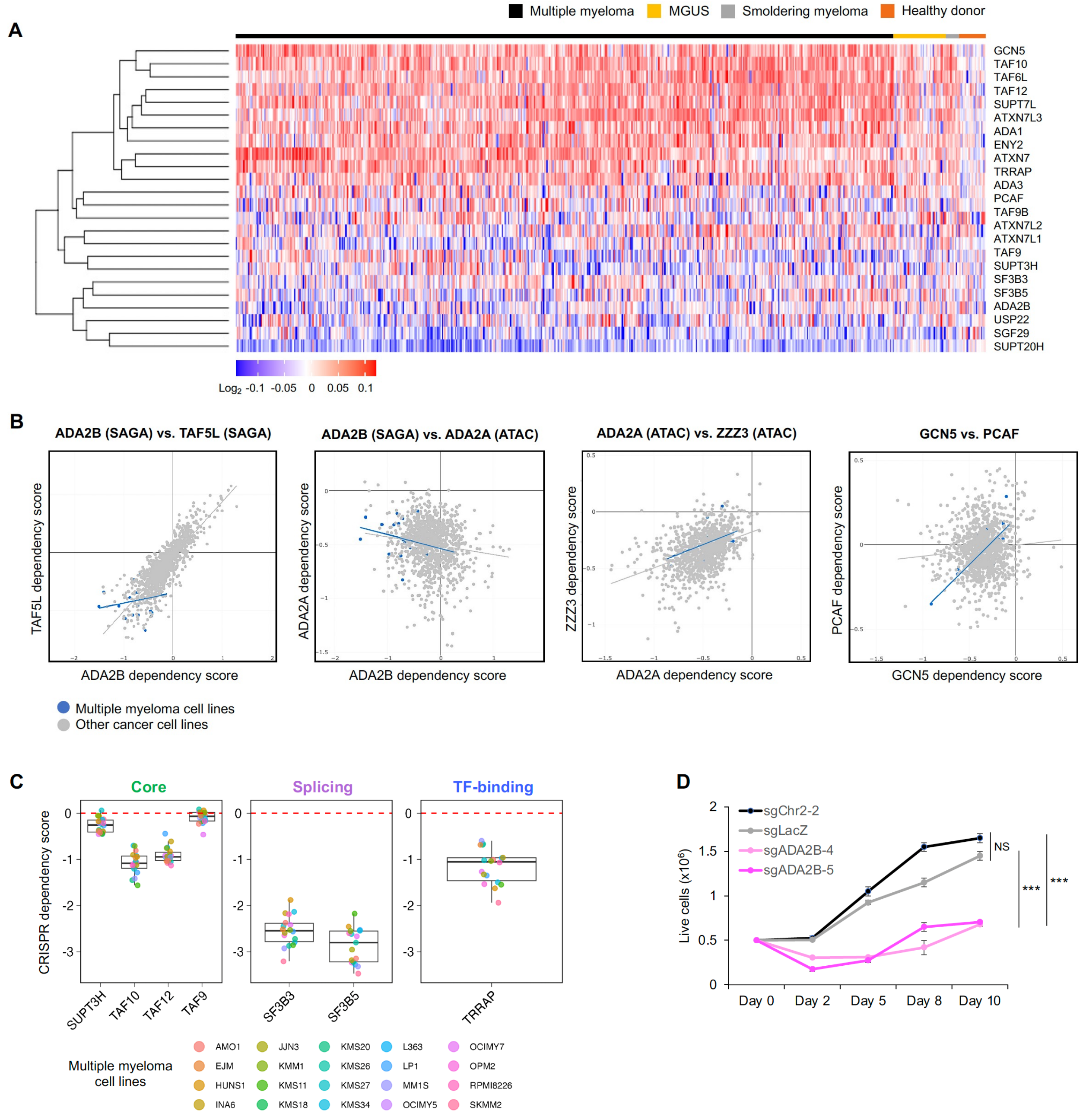


**Supplemental Figure S1. SAGA complex members are important for MM cell growth.**
**A)** Heatmap showing expression of genes encoding SAGA components in MM patients, MGUS phase, smoldering myeloma, and healthy donor plasma cells. **B)** Correlation plots of cancer dependency scores of SAGA or ATAC component genes, showing positive correlation between SAGA components, negative correlation between ADA2B (SAGA) and ADA2A (ATAC), and positive correlation between ATAC components. **C)** Dependency scores of component genes of SAGA core, splicing, and TF-binding modules for MM cell lines. **D)** Cell viability of ADA2B KO and controls sgChr2-2 and sgLacZ in MM.1S cells. A one-way Analysis of Variance (ANOVA) was performed to analyze the differences among group means, followed by the Tukey HSD post hoc test to determine whether the mean difference between specific pairs of group are statistically significant. *** $p < 0.001$, ** $p < 0.01$, * $p < 0.05$, not significant (NS) $p > 0.05$.

Supplemental Figure S2

A

| RNA-seq (FC>1.5 and FDR<0.05) | | | | | |
|---|---|---|---|---|---|
| Experiment sample | Control sample | Days of treatment | Upregulated genes | Downregulated genes | Total changed |
| shADA2B+Dox (KD) | shNT+Dox | 3 | 177 | 533 | 710 |
| shADA2B+Dox (KD) | shNT+Dox | 5 | 502 | 474 | 976 |
| shADA2B+Dox (KD) | shNT no Dox | 3 | 218 | 667 | 885 |
| shADA2B+Dox (KD) | shNT no Dox | 5 | 882 | 465 | 1347 |
| shADA2B+Dox (KD) | shADA2B no Dox | 3 | 219 | 682 | 901 |
| shADA2B+Dox (KD) | shADA2B no Dox | 5 | 620 | 507 | 1127 |
| shNT+Dox | shNT no Dox | 3 | 6 | 10 | 16 |
| shNT+Dox | shNT no Dox | 5 | 115 | 19 | 134 |
| shADA2B no Dox | shNT no Dox | 3 | 34 | 77 | 111 |
| shADA2B no Dox | shNT no Dox | 5 | 40 | 1 | 41 |

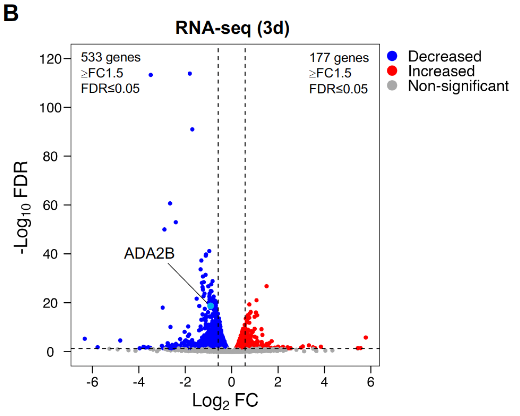


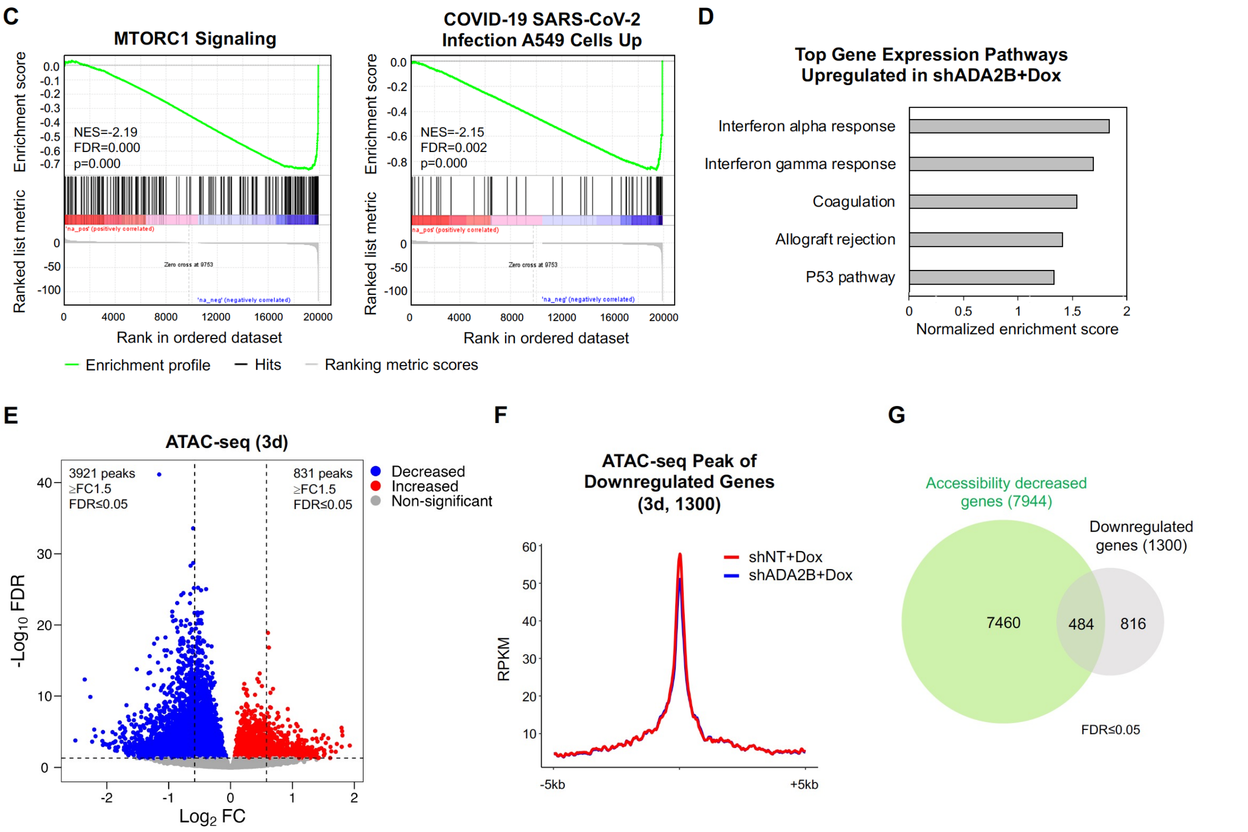


**Supplemental Figure S2. Transcriptome and chromatin accessibility changes upon ADA2B depletion.**

**A)** Table of statistically significant, markedly upregulated and downregulated genes (FC ≥ 1.5, FDR ≤ 0.05) detected by RNA-seq in each pair of comparison. Dox-induced ADA2B KD samples are shown in pink. Dox treatment for 3 days had minimal effects on the transcriptome, as indicated by the small number of differentially expressed genes; Dox treatment for 5 days caused gene upregulation but minimal downregulation. **B)** Volcano plot of gene expression changes detected by RNA-seq in shADA2B cells relative to shNT cells after 3 days of Dox treatment. Genes with statistically significant differential expression with FC ≥ 1.5 include 533 downregulated genes and 177 upregulated genes. **C)** GSEA enrichment plots of MTORC1 signaling and upregulated genes in A549 upon SARS-CoV-2 Infection (COVID-19 SARS-CoV-2 Infection A549 Cells Up, 24 hours postinfection) that were enriched in downregulated genes in shADA2B cells relative to shNT cells after 3 days of Dox treatment. **D)** Top five gene expression pathways identified by GSEA that were enriched in upregulated genes in shADA2B cells relative to shNT cells after 5 days of Dox treatment. **E)** Volcano plot of chromatin accessibility changes detected by ATAC-seq in shADA2B cells relative to shNT cells after 3 days of Dox treatment. Statistically significant differential peaks with FC ≥ 1.5 include 3921 decreased peaks and 831 increased peaks. **F)** Average RPKM (reads per kilobase per million) normalized signal of ATAC-seq peaks associated to all (1300) downregulated genes in shADA2B cells relative to shNT cells after 5 days of Dox treatment. **G)** Venn diagram depicting the overlap between downregulated genes (1300 genes) and genes with decreased accessibility (7944 genes) in shADA2B cells relative to shNT cells after 3 days of Dox treatment.

Supplemental Figure S3

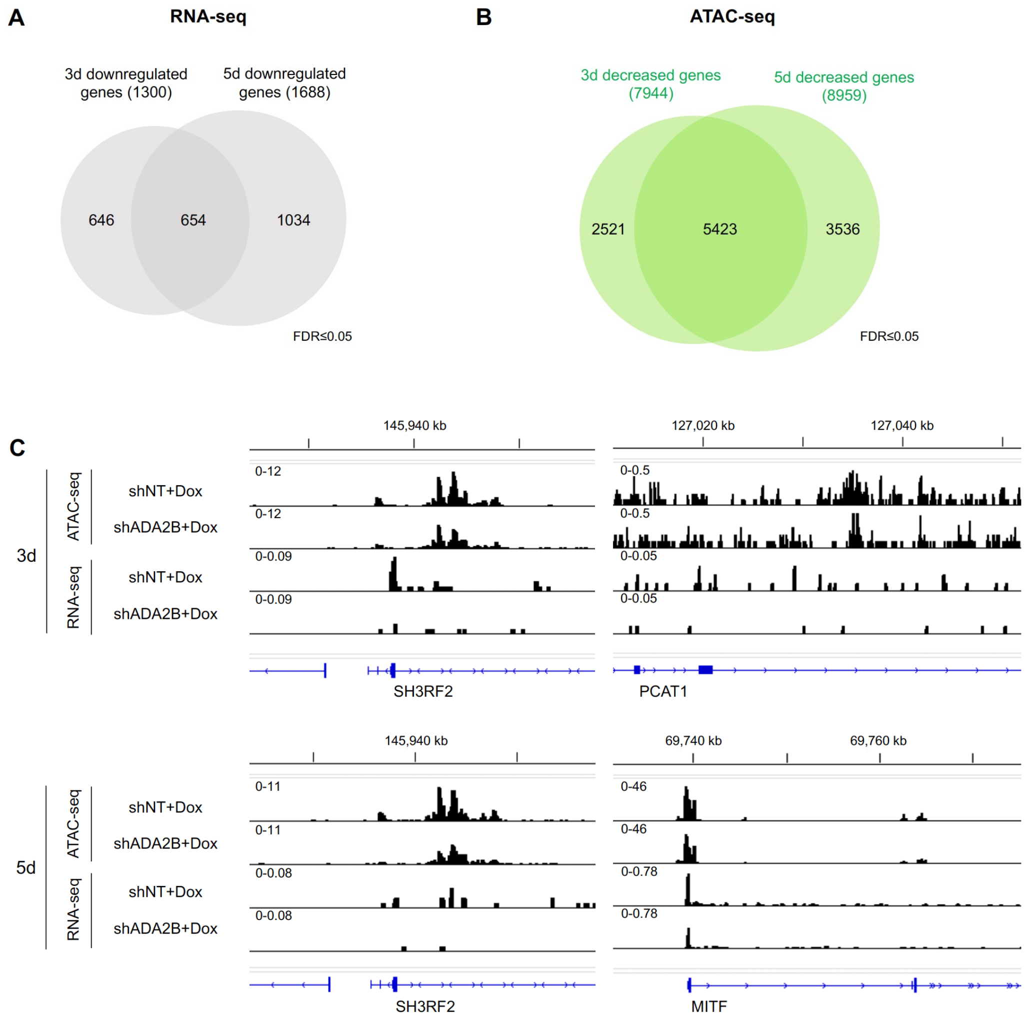


**Supplemental Figure S3. Genes with sustained decrease in expression or chromatin accessibility upon ADA2B depletion.**

**A)** Venn diagram depicting the overlap between 3-day and 5-day statistically significant downregulated genes in shADA2B cells relative to shNT cells after Dox treatment. **B)** Venn diagram depicting the overlap between 3-day and 5-day genes with statistically significant decreased accessibility in shADA2B cells relative to shNT cells after Dox treatment. **C)** RNA-seq and ATAC-seq tracks at oncogenes with decreased expression as well as accessibility in shADA2B cells relative to shNT cells after Dox treatment, including SH3RF2 (at both 3 and 5 days of treatment), PCAT1 (3 days), and MITF (5 days).

Supplemental Figure S4

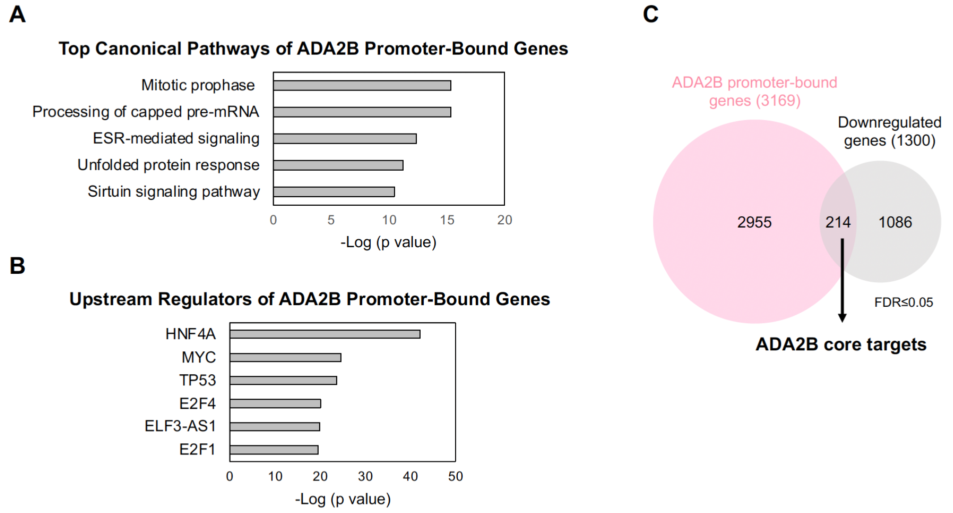


**Supplemental Figure S4. Analyses of ADA2B target genes.**
**A)** Top five canonical pathways of genes bound by ADA2B at their promoters as identified by IPA. **B)** Top five upstream regulators of genes bound by ADA2B at their promoters as identified by IPA. **C)** Venn diagram depicting the overlap between ADA2B promoter-bound genes (3169 genes) and downregulated genes (1300 genes) in shADA2B cells relative to shNT cells after 3 days of Dox treatment, identifying 214 ADA2B core targets.

Supplemental Figure S5

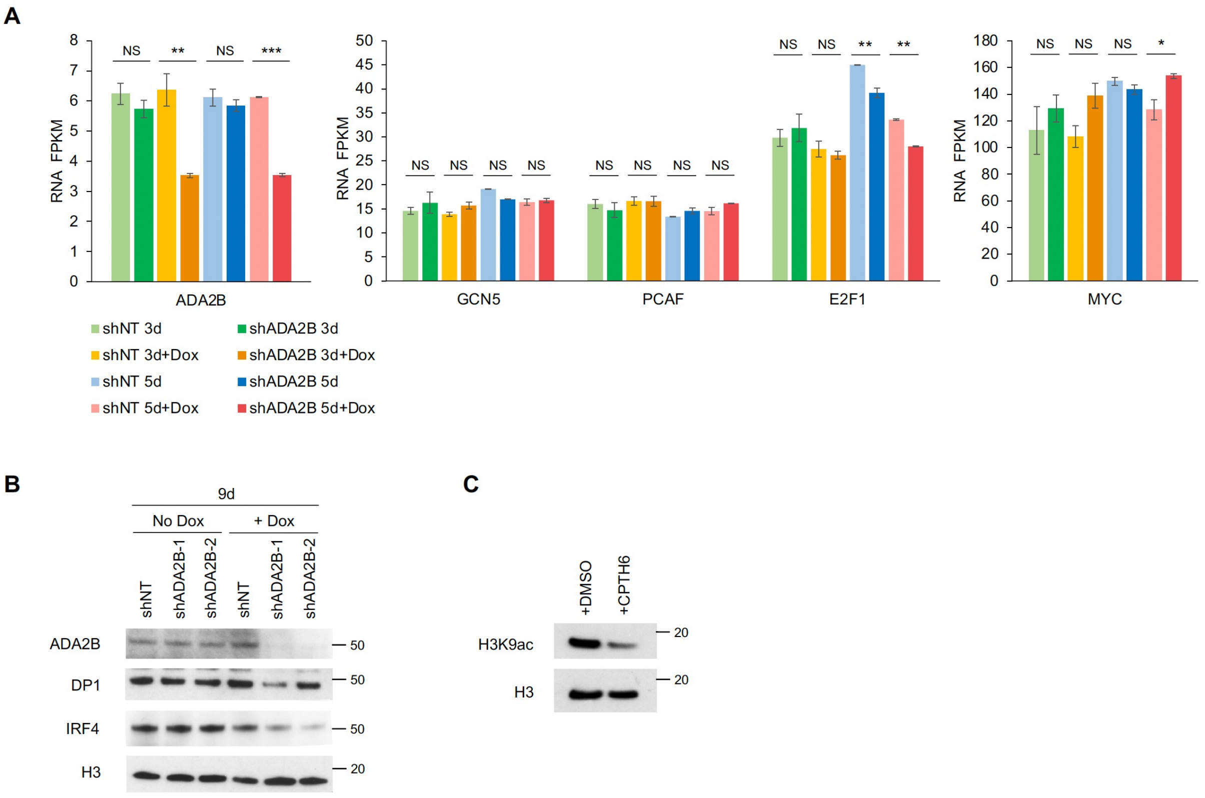


**Supplemental Figure S5. Impact of ADA2B loss on expression of transcription factors in MM.** **A)** Immunoblots showing levels of DP1 and IRF4 in shNT and shADA2B cells with and without Dox treatment for 9 days. Histone H3 is included as a loading control. **B)** Transcript levels in FPKM (fragments per kilobase per million) of *ADA2B*, *GCN5*, *PCAF*, *E2F1*, and *MYC* in shNT and shADA2B cells with and without Dox treatment for 3 and 5 days as determined by RNA-seq. **C)** Immunoblots showing global H3K9ac levels in MM.1S cells treated with 10 μM CPTH6 for 6 days in comparison to DMSO vehicle control. Histone H3 is included as a loading control. A one-way Analysis of Variance (ANOVA) was performed to analyze the differences among group means, followed by the Tukey HSD post hoc test to determine whether the mean difference between specific pairs of group are statistically significant. *** $p < 0.001$, ** $p < 0.01$, * $p < 0.05$, not significant (NS) $p > 0.05$.

Supplemental Figure S6

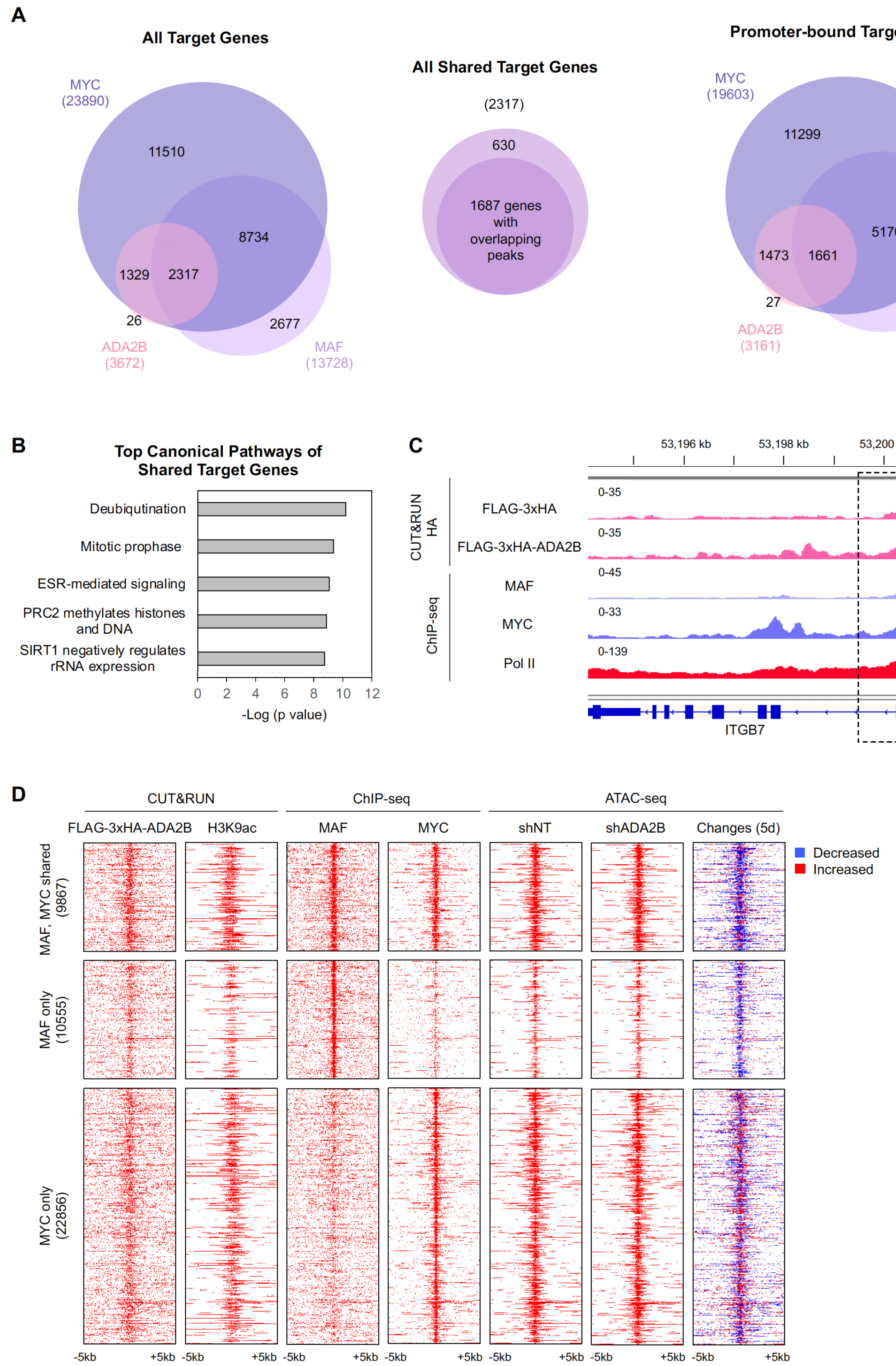


**Supplemental Figure S6. ADA2B, MAF, and MYC share target genes.**
**A)** Venn diagrams depicting the overlaps between ADA2B, MAF, and MYC all target genes and target genes bound by ADA2B, MAF, or MYC at their promoters. Among their shared target genes, number of genes with overlapping binding peaks of all three regulators are also shown. **B)** Top five canonical pathways of genes with overlapping binding sites of ADA2B, MAF, and MYC at their promoters as identified by IPA. **C)** Tracks of FLAG-3xHA-ADA2B, MAF, MYC, and Pol II binding at *ITGB7*. Regions with colocalized binding sites of FLAG-3xHA-ADA2B, MAF, and MYC are indicated. ChIP-seq tracks of MAF, MYC, and Pol II represent FC over input. **D)** Heatmaps of FLAG-3xHA-ADA2B, MAF, MYC, and ATAC-seq peaks at MAF and MYC shared binding sites and at sites bound by only MAF or MYC. FLAG-3xHA-ADA2B signals are normalized to IgG signals. ChIP-seq tracks of MAF, MYC, and Pol II represent FC over input. Changes of ATAC-seq peak signals in shADA2B cells relative to shNT cells after 5 days of Dox treatment are also shown, with decreased peaks are in blue and increased peaks in red.

Supplemental Figure S7

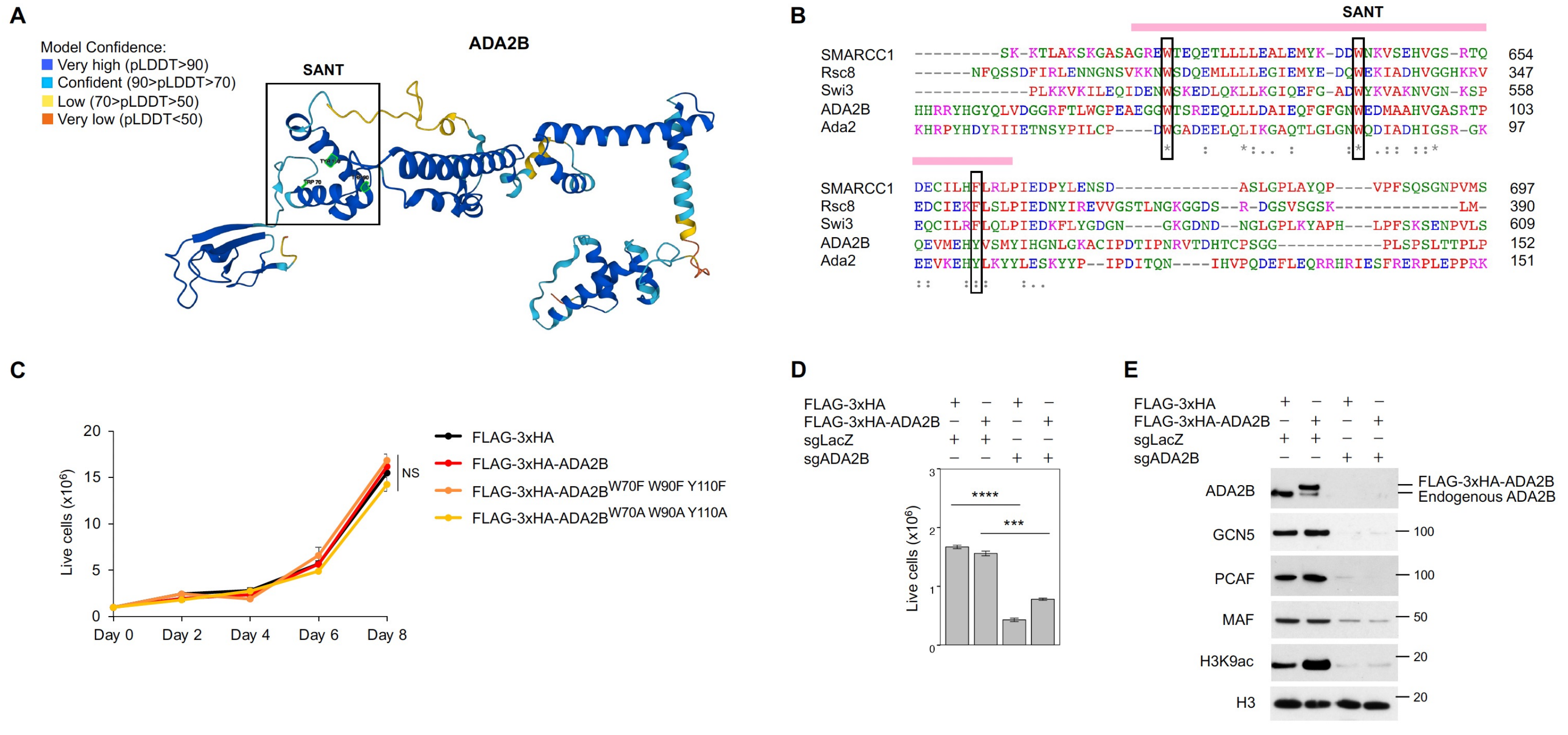


**Supplemental Figure S7. Analyses of the SANT domain of ADA2B and exogenous FLAG-3xHA-ADA2B.**
**A)** Structure of the full-length ADA2B protein as predicted by AlphaFold. The SANT domain is indicated and three predicted core hydrophobic residues (W70, W90, Y110) are highlighted in green. AlphaFold produces a per-residue confidence score (pLDDT) between 0 and 100. **B)** Sequence alignment of the SANT domain (highlighted in pink) from human ADA2B, SMARCC1, and yeast Rsc8, Swi3, Ada2 proteins by Clustal Omega. The three predicted core hydrophobic residues are indicated. **C)** Cell viability assay of MM.1S cells expressing FLAG-3xHA, FLAG-3xHA-ADA2B, FLAG-3xHA-ADA2B$^{W70A\ W90A\ Y110A}$, or FLAG-3xHA-ADA2B$^{W70F\ W90F\ Y110F}$. A one-way Analysis of Variance (ANOVA) was performed to analyze the differences among group means, followed by the Tukey HSD post hoc test to determine whether the mean difference between specific pairs of group are statistically significant. Not significant (NS) $p > 0.05$. **D)** Cell viability assay of MM.1S cells expressing FLAG-3xHA or FLAG-3xHA-ADA2B along with sgRNAs either sgLacZ or sgADA2B. Cells harboring both FLAG-3xHA-ADA2B and sgADA2B displayed significantly reduced viability compared to those harboring FLAG-3xHA-ADA2B and sgLacZ; cells expressing both FLAG-3xHA and sgADA2B showed significantly reduced viability compared to those expressing FLAG-3xHA and sgLacZ. A one-way Analysis of Variance (ANOVA) was performed to analyze the differences among group means, followed by the Tukey HSD post hoc test to determine whether the mean difference between specific pairs of group are statistically significant. **** $p < 0.0001$, *** $p < 0.001$, ** $p < 0.01$, * $p < 0.05$, not significant (NS) $p > 0.05$. **E)** Immunoblots showing depletion of FLAG-3xHA-ADA2B, endogenous ADA2B, GCN5, PCAF, MAF, and global H3K9ac in the two cell lines displaying reduced viability in panel D. Histone H3 blot is included as a loading control.
